# The *Drosophila* SWI/SNF chromatin-remodeling complexes BAP and PBAP play separate roles in regulating growth and cell fate during regeneration

**DOI:** 10.1101/326439

**Authors:** Yuan Tian, Rachel K. Smith-Bolton

## Abstract

To regenerate, damaged tissue must heal the wound, regrow to the proper size, replace the correct cell types, and return to the normal gene-expression program. However, the mechanisms that temporally and spatially control the activation or repression of important genes during regeneration are not fully understood. To determine the role that chromatin modifiers play in regulating gene expression after tissue damage, we induced ablation in *Drosophila* imaginal wing discs, and screened for chromatin regulators that are required for epithelial tissue regeneration. Here we show that many of these genes are indeed important for promoting or constraining regeneration. Specifically, the two SWI/SNF chromatin-remodeling complexes play distinct roles in regulating different aspects of regeneration. The PBAP complex regulates regenerative growth and developmental timing, and is required for the expression of JNK signaling targets and the growth promoter *Myc*. By contrast, the BAP complex ensures correct patterning and cell fate by stabilizing expression of the posterior gene *engrailed*. Thus, both SWI/SNF complexes are essential for proper gene expression during tissue regeneration, but they play distinct roles in regulating growth and cell fate.

**Summary statement:** During regeneration of the *Drosophila* wing disc, the SWI/SNF PBAP complex is required for regenerative growth and expression of JNK signaling targets, while the BAP complex maintains posterior cell fate.

## Introduction

Regeneration is a complex yet highly elegant process that some organisms can use to recognize, repair and replace missing or damaged tissue. Imaginal disc repair in *Drosophila* is a good model system for understanding regeneration due to the high capacity of these tissues to regrow and restore complex patterning, as well as the genetic tools available in this model organism (Hariharan and Serras, 2017). Regeneration requires the coordinated expression of genes that regulate the sensing of tissue damage, induction of regenerative growth, repatterning of the tissue, and coordination of regeneration with developmental timing. Initiation of regeneration in imaginal discs requires known signaling pathways such as the ROS, JNK, Wg, p38, Jak/STAT, and Hippo pathways (Bergantinos et al., 2010; Bosch et al., 2008; Grusche et al., 2011; Katsuyama et al., 2015; Santabárbara-Ruiz et al., 2015; Schubiger et al., 2010; Smith-Bolton et al., 2009; Sun and Irvine, 2011). These pathways activate many regeneration genes, such as the growth promoter Myc (Smith-Bolton et al., 2009) and the hormone-like peptide *ilp8*, which delays pupariation after imaginal disc damage (Colombani et al., 2012; Garelli et al., 2012). However, misregulation of these signals can impair regeneration. For example, elevated levels of JNK signaling can induce patterning defects in the posterior of the wing (Schuster and Smith-Bolton, 2015), and elevated ROS levels can suppress JNK activity and regenerative growth (Brock et al., 2017). While the signals that initiate regeneration have been extensively studied, regulation of regeneration gene expression in response to tissue damage is not fully understood.

Such regulation could occur through chromatin modification. In *Drosophila*, chromatin modifiers include the repressive complexes PRC1 and PRC2, the activating complexes TAC1, COMPASS and COMPASS-like, and the SWI/SNF chromatin remodelers BAP and PBAP (Kassis et al., 2017). PRC2 carries out trimethylation of histone H3 at lysine 27, recruiting PRC1 to repress transcription of nearby genes. COMPASS-like and COMPASS carry out histone H3 lysine 4 monomethylation and di- and trimethylation, respectively, thereby activating expression of nearby genes. TAC1 acetylates histone H3 lysine 27, also supporting activation of gene transcription. BAP and PBAP alter or move nucleosomes to facilitate binding of transcription factors and chromatin modifiers. Rapid changes in gene expression induced by these complexes may help facilitate a damaged tissue’s regenerative response.

A few chromatin modifiers and histone modifications have been reported to be important for regulating regeneration of Xenopus tadpole tails, mouse pancreas and liver, zebrafish fins, and *Drosophila* imaginal discs (Blanco et al., 2010; Fukuda et al., 2012; Jin et al., 2015; Pfefferli et al., 2014; Scimone et al., 2010; Skinner et al., 2015; Stewart et al., 2009; Tseng et al., 2011; Wang et al., 2008). Furthermore, components of *Drosophila* and mouse SWI/SNF complexes regulate regeneration in the *Drosophila* midgut and mouse skin, liver, and ear (Jin et al., 2013; Sun et al., 2016; Xiong et al., 2013). However, little is known about how these complexes alter gene expression, signaling, and cellular behavior to regulate regeneration. Importantly, genome-wide analysis of chromatin state after *Drosophila* imaginal disc damage revealed changes in chromatin around a large set of genes, including known regeneration genes (Vizcaya-Molina et al., 2018). Thus, chromatin modifiers likely play a key role in regulating activation of the regeneration program. However, it is unclear whether all regeneration genes are coordinately regulated in the same manner, or whether specific chromatin modification complexes target different subsets of genes that respond to tissue damage.

To probe the role of chromatin modifiers in tissue regeneration systematically, we assembled a collection of pre-existing *Drosophila* mutants and RNAi lines targeting components of these complexes as well as other genes that regulate chromatin, and screened these lines for regeneration defects using the *Drosophila* wing imaginal disc. We used a spatially and temporally controllable tissue-ablation method that uses transgenic tools to induce tissue damage only in the wing primordium (Smith-Bolton et al., 2009). This method ablates 94% of the wing primordium on average at the early third instar and allows the damaged wing discs to regenerate *in situ*. Previous genetic screens using this tissue ablation method have identified genes critical for regulating different aspects of regeneration, such as *taranis*, *trithorax*, and *cap-n-collar*, demonstrating its efficacy in finding regeneration genes (Brock et al., 2017; Schuster and Smith-Bolton, 2015; Skinner et al., 2015).

Through this targeted genetic screen of chromatin regulators we found that mutations in *Drosophila* SWI/SNF components caused striking regeneration defects. The SWI/SNF complexes are conserved multi-subunit protein complexes that activate or repress gene expression (Wilson and Roberts, 2011) by using the energy from ATP hydrolysis to disrupt histone-DNA contacts and remodel nucleosome structure and position (Côté et al., 1994; Kwon et al., 1994). Brm is the only ATPase of the SWI/SNF complexes in *Drosophila* (Kassis et al., 2017; Tamkun et al., 1992). Moira (Mor) serves as the core scaffold of the complexes (Mashtalir et al., 2018). Other components contain domains involved in protein-protein interactions, protein-DNA interactions, or interactions with modified histones (Hargreaves and Crabtree, 2011). There are two subtypes of SWI/SNF in *Drosophila*: the Brahma-associated proteins (BAP) and the Polybromo-associated BAP (PBAP) remodeling complexes (Collins and Treisman, 2000; Mohrmann et al., 2004). They share common core components, including Brm, Snr1, Mor, Bap55, Bap60, Bap111 and Actin (Mohrmann et al., 2004), but contain different signature proteins. The PBAP complex is defined by the components Bap170, Polybromo and Sayp (Mohrmann et al., 2004; Chalkley et al., 2008). Osa defines the BAP complex (Collins et al., 1999; Vázquez et al., 1999).

Here we show that the SWI/SNF complexes BAP and PBAP are required for regeneration, and that the two complexes play distinct roles. The PBAP complex is important for activation of JNK signaling targets such as *ilp8* to delay metamorphosis and allow enough time for the damaged tissue to regrow, and for expression of *myc* to drive regenerative growth. By contrast, the BAP complex is not required for regenerative growth, but instead functions to prevent changes in cell fate induced by tissue damage through stabilizing expression of the posterior identity gene *engrailed*. Thus, different aspects of the regeneration program are regulated independently by distinct chromatin regulators.

## Materials and Methods

### Fly stocks

The following fly stocks were obtained for this study. In some cases they were rebalanced before performing experiments: *w*^*1118*^*;; rnGAL4, UAS-rpr, tub-GAL80*^*ts*^*/TM6B, tubGAL80* (Smith-Bolton et al., 2009), *w*^*1118*^ (Wild type), *w*; P{neoFRT}82B osa*^*308*^*/TM6B, Tb*^*1*^ (Bloomington *Drosophila* stock center, BL#5949) (Treisman et al., 1997), *w*; Bap170*^*Δ135*^*/T(2;3)SM6a-TM6B, Tb*^*1*^ was a gift from Jessica E. Treisman (Carrera et al., 2008), *brm*^*2*^ *e*^*s*^ *ca*^*1*^*/TM6B, Sb*^*1*^ *Tb*^*1*^ *ca*^*1*^ (BL#3619) (Kennison and Tamkun, 1988), *mor*^*1*^*/TM6B, Tb*^*1*^ (BL#3615) (Kennison and Tamkun, 1988), *y*^*1*^ *w*^*1*^*; P{neoFRT}40A P{FRT(w*^*hs*^*)}G13 cn*^*1*^ *PBac{SAs-topDsRed}Bap55*^*LL05955*^ *bw*^*1*^*/CyO, bw*^*1*^ (BL#34495) (Schuldiner et al., 2008), *bap111* RNAi (Vienna *Drosophila* Resource Center, VDRC#104361), control RNAi background (VDRC#15293) *bap60* RNAi (VDRC#12673), *brm* RNAi (VDRC#37721), *P{PZ}tara*^*03881*^ *ry*^*506*^*/TM3, ry*^*RK*^ *Sb*^*1*^ *Ser*^*1*^ (BL#11613) (Gutierrez, 2003), *UAS-tara* was a gift from Michael Cleary (Manansala et al., 2013), *TRE-Red* was a gift from Dirk Bohmann (Chatterjee and Bohmann, 2012). *mor*^*2*^, *mor*^*11*^ and *mor*^*12*^ alleles were gifts from James Kennison (Kennison and Tamkun, 1988), *snr1*^*E2*^ and *snr1*^*SR21*^ alleles were gifts from Andrew Dingwall (Zraly et al., 2003).

The mutants and RNA interference lines in Table S1 used for the chromatin regulator screen were:

*st*^*1*^ *in*^*1*^ *kni*^*ri-1*^ *Scr*^*W*^ *Pc*^*3*^*/TM3, Sb*^*1*^ *Ser*^*1*^ (BL#3399),
*cn*^*1*^ *Psc*^*1*^ *bw*^*1*^ *sp*^*1*^*/CyO* (BL#4200),
*y*^*1*^ *w*; P{neoFRT}42D Psc*^*e24*^*/SM6b, P{eve-lacZ8.0}SB1* (BL#24155),
*w*; P{neoFRT}82B Abd-B*^*Mcp-1*^ *Sce*^*1*^*/TM6C, Sb*^*1*^ *Tb*^*1*^ (BL#24618),
*w*; P{neoFRT}82B Scm*^*D1*^*/TM6C, Sb*^*1*^ *Tb*^*1*^ (BL#24158),
*w*; E(z)*^*731*^ *P{1xFRT.G}2A/TM6C, Sb*^*1*^ *Tb*^*1*^ (BL#24470),
*w*; Su(z)12*^*2*^ *P{FRT(w*^*hs*^*)}2A/TM6C, Sb*^*1*^ *Tb*^*1*^ (BL#24159),
*esc*^*21*^ *b*^*1*^ *cn*^*1*^*/In(2LR)Gla, wg*^*Gla-1*^*; ca*^*1*^ *awd*^*K*^ (BL#3623),
*y*^*1*^ *w*^*67c23*^*; P{wHy}Caf1-55*^*DG25308*^ (BL#21275),
*w*^*1118*^*; P{XP}escl*^*d01514*^ (BL#19163),
*y*^*1*^ *w*; phol*^*81A*^*/TM3, Ser*^*1*^ *y*^*+*^ (BL#24164),
*red*^*1*^ *e*^*1*^ *ash2*^*1*^*/TM6B, Tb*^*1*^ (BL#4584),
*w*^*1118*^*; PBac{WH}Utx*^*f01321*^*/CyO* (BL#18425),
*w*; ash1*^*22*^ *P{FRT(w*^*hs*^*)}2A/TM6C, Sb*^*1*^ *Tb*^*1*^ (BL#24161),
*w*^*1118*^*; E(bx)*^*Nurf301-3*^*/TM3, P{ActGFP}JMR2, Ser*^*1*^ (BL#9687),
*y*^*1*^ *w*^*67c23*^*; P{lacW}Nurf-38*^*k16102*^*/CyO* (BL#12206),
*Mi-2*^*4*^ *red*^*1*^ *e*^*4*^*/TM6B, Sb*^*1*^ *Tb*^*1*^ *ca*^*1*^ (BL#26170),
*mor* RNAi (VDRC#6969),
*psq*^*E39*^*/CyO; ry*^*506*^ (BL#7321),
*Rbf*^*14*^ *w*^*1118*^*/FM7c* (BL#7435),
*w*^*1118*^ *P{EP}Dsp1*^*EP355*^ (BL#17270),
*cn*^*1*^ *grh*^*IM*^ *bw*^*1*^*/SM6a* (BL#3270),
*y*^*1*^ *w*^*67c23*^*; P{lacW}lolal*^*k02512*^*/CyO* (BL#10515),
*w*; P{neoFRT}42D Pcl*^*5*^*/CyO* (BL#24157),
*w*; HDAC1*^*def24*^ *P{FRT(w*^*hs*^*)}2A P{neoFRT}82B/TM6B, Tb*^*1*^ (BL#32239),
*w*^*1118*^*; Sirt1*^*2A-7-11*^ (BL#8838),
*Eip74EF*^*v4*^ *vtd*^*4*^*/TM3, st*^*24*^ *Sb*^*1*^ (BL#5050),
*sc*^*1*^ *z*^*1*^ *w*^*is*^*; Su(z)2*^*1.b7*^*/CyO* (BL#5572),
*P{PZ}gpp*^*03342*^ *ry*^*506*^*/TM3, ry*^*RK*^ *Sb*^*1*^ *Ser*^*1*^ (BL#11585),
*y*^*1*^ *w*^*1118*^*; P{lacW}mod(mdg4)*^*L3101*^*/TM3, Ser*^*1*^ (BL#10312),
*w*^*1118*^*; PBac{RB}su(Hw)*^*e04061*^*/TM6B, Tb*^*1*^ (BL#18224),
*cn*^*1*^ *P{PZ}lid*^*10424*^*/CyO; ry*^*506*^ (BL#12367),
*Asx*^*XF23*^*/CyO* (BL#6041),
*y*^*1*^ *w*^*1*^*; P{neoFRT}40A P{FRT(w*^*hs*^*)}G13 cn*^*1*^ *PBac{SAsto-pDsRed}dom*^*LL05537*^ *bw*^*1*^*/CyO, bw*^*1*^ (BL#34496),
*cn*^*1*^ *E(Pc)*^*1*^ *bw*^*1*^*/SM5* (BL#3056),
*kis*^*1*^ *cn*^*1*^ *bw*^*1*^ *sp*^*1*^*/SM6a* (BL#431),
*kto*^*1*^ *ca*^*1*^*/TM6B, Tb*^*1*^ (BL#3618),
*skd*^*2*^*/TM6C, cu*^*1*^ *Sb*^*1*^ *ca*^*1*^ (BL#5047).

### Genetic screen

Mutants or RNAi lines were crossed to *w*^*1118*^*;; rnGAL4, UAS-rpr, tub-GAL80*^*ts*^*/TM6B, tubGAL80* flies. Controls were *w*^*1118*^ or the appropriate RNAi background line. Embryos were collected at room temperature on grape plates for 4 hours in the dark, then kept at 18°C. Larvae were picked at 2 days after egg lay into standard Bloomington cornmeal media and kept at 18°C, 50 larvae in each vial, 3 vials per genotype per replicate. On day 7, tissue ablation was induced by a placing the vials in a 30°C circulating water bath for 24 hours. Then ablation was stopped by placing the vials in ice water for 60 seconds and returning them to 18°C for regeneration. The regeneration index was calculated by summing the product of approximate wing size (0%, 25%, 50%, 75% and 100%) and the corresponding percentage of wings for each wing size. The Δ Index was calculated by subtracting the regeneration index of the control from the regeneration index of the mutant or RNAi line.

To observe and quantify the patterning features and absolute wing size, adult wings that were 75% size or greater were mounted in Gary’s Magic Mount (Canada balsam (Sigma) dissolved in methyl salicylate (Sigma)). The mounted adult wings were imaged with an Olympus SZX10 microscope using an Olympus DP21 camera, with the Olympus CellSens Dimension software. Wings were measured using ImageJ.

### Immunostaining

Immunostaining was carried out as previously described (Smith-Bolton et al., 2009). Primary antibodies used in this study were rabbit anti-Myc (1:500; Santa Cruz Biotechnology), mouse anti-Nubbin (1:250; gift from Steve Cohen) (Ng et al., 1996), mouse anti-engrailed/invected (1:3; Developmental Studies Hybridoma Bank (DSHB)) (Patel et al., 1989), mouse anti-Patched (1:50; DSHB) (Capdevila et al., 1994), mouse anti-Achaete (1:10; DSHB) (Skeath and Carroll, 1992), rabbit anti-PH3 (1:500; Millipore), mouse anti-Osa (1:1; DSHB) (Treisman et al., 1997), rat anti-Ci (1:10; DSHB) (Motzny and Holmgren, 1995), rabbit anti-Dcp1 (1:250; Cell Signaling), mouse anti-βgal (1:100; DSHB), rabbit anti-phos-pho-Mad (1:100; Cell Signaling), mouse anti-Mmp1 (1:10 of 1:1:1 mixture of monoclonal antibodies 3B8D12, 5H7B11, and 3A6B4, DSHB)(Page-McCaw et al., 2003). The Developmental Studies Hybridoma Bank (DSHB) was created by the NICHD of the NIH and is maintained at the University of Iowa, Department of Biology, Iowa City, IA 52242. Secondary antibodies used in this study were AlexaFluor secondary antibodies (Molecular Probes) (1:1000). TO-PRO-3 iodide (Molecular Probes) was used to detect DNA at 1:500.

Confocal images were collected with a Zeiss LSM700 Confocal Microscope using ZEN software (Zeiss). Images were processed with ImageJ (NIH) and Photoshop (Adobe). Average fluorescence intensity was measured by ImageJ. Quantification of fluorescence intensity and phospho-histone H3 positive cells was restricted to the wing pouch, as marked by anti-Nubbin immunostaining or morphology. The area of the regenerating wing primordium was quantified by measuring the anti-Nubbin immunostained area in ImageJ.

### Quantitative RT-PCR

qPCR was conducted as previously described (Skinner et al., 2015). Each independent sample consisted of 50 wing discs. 3 biological replicates were collected for each genotype and time point. Expression levels were normalized to the control *gapdh2*. The fold changes compared to the *w*^*1118*^ undamaged wing discs are shown. Primers used in the study were:

*GAPDH2* (Forward: 5’-GTGAAGCTGATCTCTTGGTACGAC-3’; Reverse: 5’-CCGCGCCCTAATCTTTAACTTTTAC-3’),
*ilp8* (Qiagen QT00510552),
*mmp1* (Forward: 5’-TCGGCTGCAAGAACACGCCC-3’; Reverse: 5’-CGCCCACGGCTGCGTCAAAG-3’),
*moira* (Forward: 5’-GATGAGGTGCCCGCTACAAT-3’; Reverse: 5’-CTGCTGCGGTTTCGTCTTTT-3’),
*brm* (Forward: 5’-GCACCACCAGGGGATGATTT-3’; Reverse: 5’-TTGTGTGGGTGCATTGGGT-3’),
*Bap60* (Forward: 5’-AGACGAGGGATTTGAAGCTGA-3’; Reverse: 5’-AGGTCTCTTGACGGTGGACT-3’)
*myc* (Forward: 5’-CGATCGCAGACGACAGATAA-3’; Reverse: 5’-GGGCGGTATTAAATGGACCT-3’)

### Pupariation timing experiments

To quantify the pupariation rates, pupal cases on the side of each vial were counted at 24-hour intervals starting from the end of tissue ablation until no new pupal cases formed. Three independent biological replicates, which consisted of 3 vials each with 50 animals per vial, were performed for each experiment. The median day is the day on which ≥ 50% of the animals had pupariated.

### Data Availability

All relevant data are available at databank.illinois.edu.XXXXXXXX and upon request.

## Results

### A genetic screen of chromatin modifier mutants and RNAi lines

To identify regeneration genes among *Drosophila* chromatin regulators, we conducted a genetic screen similar to our previously reported unbiased genetic screen for genes that regulate wing imaginal disc regeneration (Brock et al., 2017)(Fig. 1A). To induce tissue ablation, *rotund-GAL4* drove the expression of the pro-apoptotic gene *UAS-reaper* in the imaginal wing pouch, and *tubulin-GAL80*^*ts*^ provided temporal control, enabling us to turn ablation on and off by varying the temperature (Smith-Bolton et al., 2009). The ablation was carried out for 24 hours during the early third instar. We characterized the quality of regeneration by assessing the adult wing size semi-quantitatively and 1) recording the numbers of wings that were 0%, 25%, 50%, 75% or 100% the length of a normal adult wing (Fig. 1A,B), and 2) identifying patterning defects by scoring ectopic or missing features. This semi-quantitative evaluation method enabled a quick screen, at a rate of 6 genotypes per week including around 1400 adult wings, and identification of both enhancers and suppressors of regeneration (Fig. 1B-E). While control animals regenerated to varying degrees depending on the extent they delayed metamorphosis in response to damage (Khan et al., 2017; Smith-Bolton et al., 2009) as well as seasonal differences in humidity and food quality (Skinner et al., 2015), the differences between the regenerative capacity of mutants and controls were consistent (Brock et al., 2017; Khan et al., 2017; Smith-Bolton et al., 2009).

**Fig 1.**
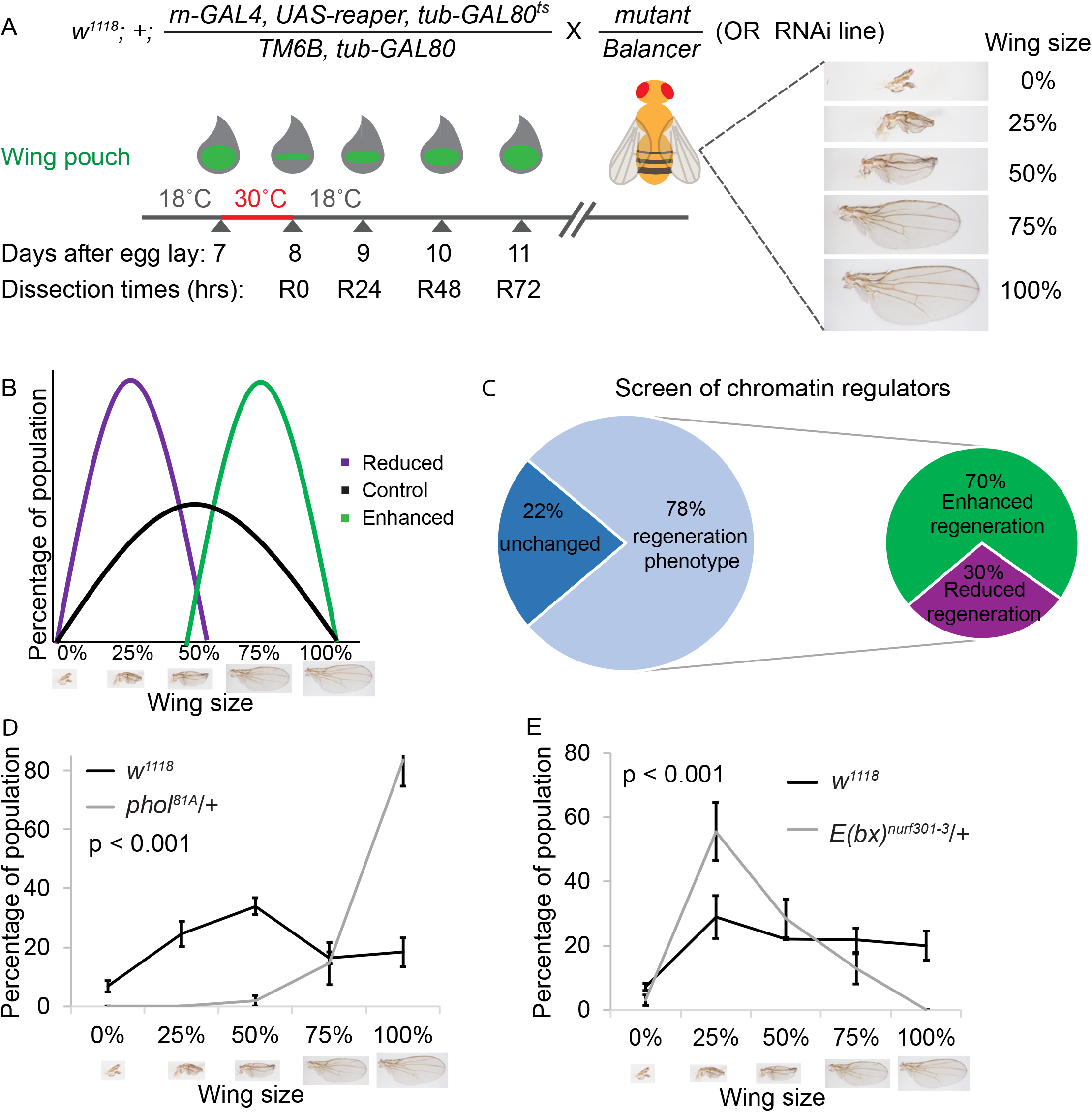
A genetic screen of chromatin regulators identified important regeneration genes. (A) Method for screening mutants or RNAi lines using a genetic ablation system. Mutants or RNAi lines of genes involved in regulating chromatin were crossed to the ablation stock (*w*^*1118*^*; +; rn-GAL4, UAS-rpr, tubGAL80*^*ts*^*/TM6B, tubGAL80*). Animals were kept at 18°C until 7 days after egg lay (AEL), when they were moved to 30°C to induce tissue ablation for 24 hours, then transferred back to 18°C to enable recovery (R). The size of the regenerated adult wings was assessed semi-quantitatively by counting the number of wings that were approximately 0%, 25%, 50%, 75% or 100% of the length of a control adult wing that had not undergone damage during the larval phase. The regenerating discs were also examined at different times denoted by hours after the beginning of recovery, such as R0, R24, R48 and R72. (B) Conceptual model for the screen to identify mutants or RNAi lines showing enhanced (green) or reduced (purple) regeneration compared to control. (C) Summary of the screen of chromatin regulators, showing percent of lines tested that had a regeneration phenotype, as well as percent of those with a phenotype that regenerated better (Δ Index ≥ 10%) or worse (Δ Index ≤ −10%) compared to controls. (D) Comparison of the size of adult wings after imaginal disc damage and regeneration in *phol*^*81A*^*/+* and wild-type (*w*^*1118*^) animals. n = 64 wings (*phol*^*81A*^*/+*) and 242 wings (*w*^*1118*^) from 3 independent experiments. Chi-square test p < 0.001 across all wing sizes. Error bars are s.e.m. (E) Comparison of the size of adult wings after imaginal disc damage and regeneration in *E(bx)*^*nurf301-3*^/+ and wild-type (*w*^*1118*^) animals. n = 219 wings (*E(bx)*^*nurf301-3*^/+) and 295 wings (*w*^*1118*^) from 3 independent experiments. Chi-square test p < 0.001 across all wing sizes. Error bars are s.e.m.

Using this system, we screened mutants and RNAi lines affecting chromatin regulators (Table S1, Fig. 1C, Fig. S1A). For each line, we calculated the Δ regeneration index, which is the difference between the regeneration indices of the line being tested and the control tested simultaneously (see materials and methods for regeneration index calculation). We set a cutoff Δ index of 10%, over which we considered the regenerative capacity to be affected. Seventy-eight percent of the mutants and RNAi lines tested had a change in regeneration index of 10% or more compared to controls (Table S1, Fig. 1C, Fig. S1A), consistent with the idea that changes in chromatin structure are required for the damaged tissue to execute the regeneration program. Twenty-two percent of the mutants and RNAi lines failed to meet our cutoff and were not pursued further (Table S1, Fig. 1C). Strikingly, 41% of the tested lines, such as *phol*^*81A*^*/+*, which affects the PhoRC complex, had larger adult wings after ablation and regeneration compared to control *w*^*1118*^ animals that had also regenerated (Fig. 1D), indicating enhanced regeneration, although none were larger than a normal-sized wing. By contrast, 25% of the tested lines, such as *E(bx)*^*nurf301-3*^*/+*, which affects the NURF complex, had smaller wings (Fig. 1E), indicating worse regeneration. Unexpectedly, mutations that affected the same complex did not have consistent phenotypes (Table S1), suggesting that chromatin modification and remodeling likely regulate a delicate balance of genes that promote and constrain regeneration. Indeed, transcriptional profiling has identified a subset of genes that are upregulated after wing disc ablation (Khan et al., 2017), some of which promote regeneration, and some of which constrain regeneration, indicating that gene regulation after tissue damage is not as simple as turning on genes that promote regeneration and turning off genes that inhibit regeneration.

### The SWI/SNF PBAP and BAP complexes have opposite phenotypes

To clarify the roles of one type of chromatin-regulating complex in regeneration, we focused on the SWI/SNF chromatin-remodeling complexes (Fig. 2A). As shown in Table S1, different components of the SWI/SNF complexes showed different phenotypes after ablation and regeneration of the wing pouches. Animals heterozygous mutant for the PBAP-specific components Bap170 (*Bap170*^*Δ135*^/+) and Polybromo (*polybromo*^*Δ86*^/+) had adult wings that were smaller after disc regeneration than *w*^*1118*^ adult wings after disc regeneration (Fig. 2B,C), suggesting that the PBAP complex is required for ablated wing discs to regrow. To confirm these semiquantitative results, we mounted adult wings and measured absolute wing sizes (N≥100 wings for each genotype). The reduced regeneration of *Bap170*^*Δ135*^/+ wing discs was confirmed by measurement of the adult wings (Fig. 2E). By contrast, animals heterozygous mutant for the BAP-specific component Osa (*osa*^*308*^/+) had larger adult wings after disc regeneration compared to *w*^*1118*^ adult wings after disc regeneration (Fig. 2D), suggesting that impairment of the BAP complex deregulates growth after tissue damage. Measurement of the adult wings of *osa*^*308*^/+ animals after disc regeneration confirmed the enhanced regeneration (Fig. 2F).

**Fig 2.**
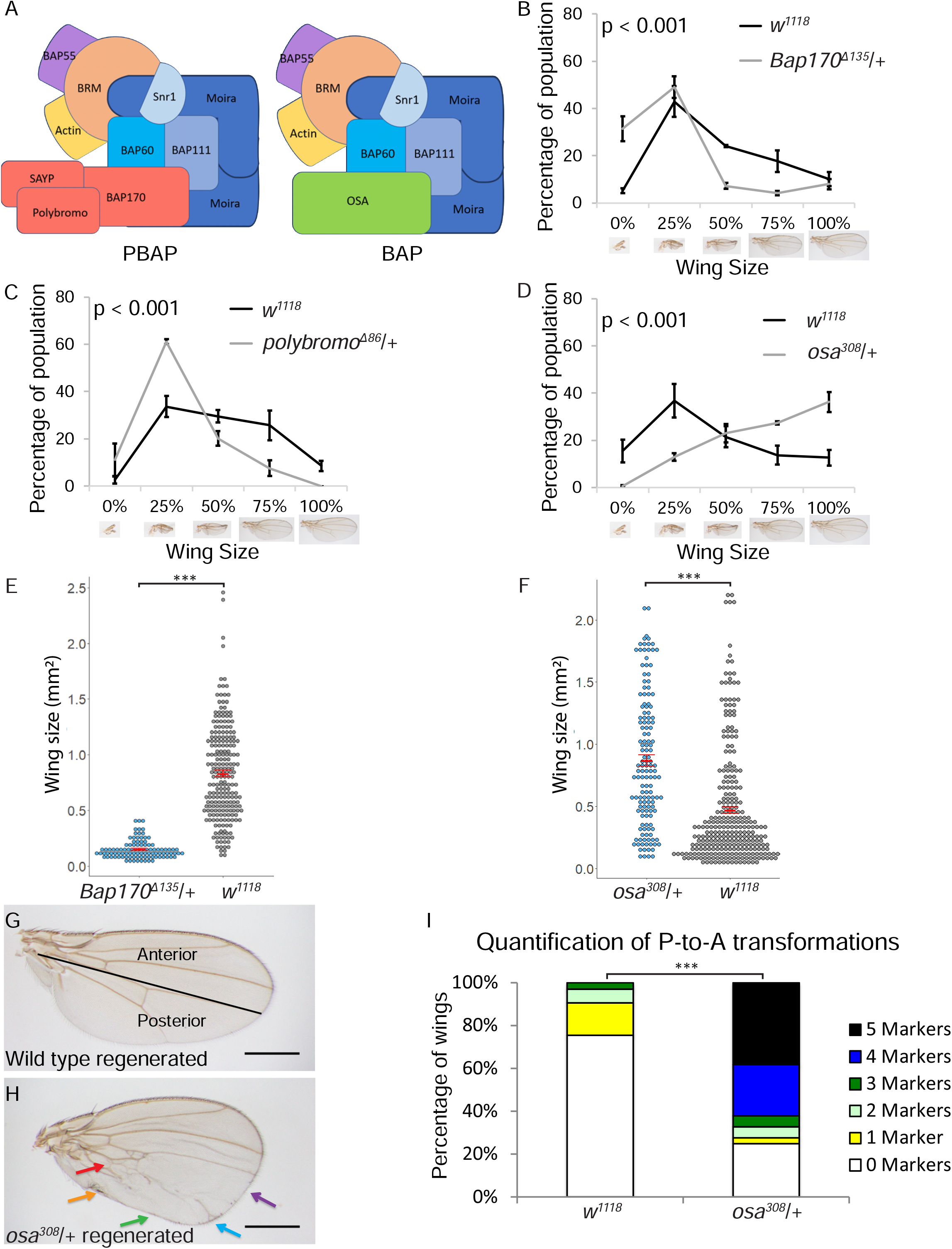
SWI/SNF components Bap170, Polybromo and Osa are required for regeneration. (A) Schematics of the two *Drosophila* SWI/SNF chromatin-remodeling complexes: BAP and PBAP, drawn based on complex organization determined in (Mashtalir et al., 2018). (B) Comparison of the size of adult wings after imaginal disc damage and regeneration in *Bap170*^*Δ135*^/+ and wild-type (*w*^*1118*^) animals. n = 190 wings (*Bap170*^*Δ135*^/+) and 406 wings (*w*^*1118*^) from 3 independent experiments. Chi-square test p < 0.001 across all wing sizes. (C) Comparison of the size of adult wings after imaginal disc damage and regeneration in *polybromo*^*Δ86*^/+ and wild-type (*w*^*1118*^) animals. n = 180 wings (*polybromo*^*Δ86*^/+) and 396 wings (*w*^*1118*^) from 3 independent experiments. Chi-square test p < 0.001 across all wing sizes. (D) Comparison of the size of adult wings after imaginal disc damage and regeneration in *osa*^*308*^/+ and wild-type (*w*^*1118*^) animals. n = 146 wings (*osa*^*308*^/+) and 296 wings (*w*^*1118*^) from three independent experiments. Chi-square test p < 0.001 across all wing sizes. (E) Wings were mounted, imaged, and measured after imaginal disc damage and regeneration in *Bap170*^*Δ135*^/+ and wild-type (*w*^*1118*^) animals. n = 100 wings (*Bap170*^*Δ135*^/+) and 224 wings (*w*^*1118*^) from 3 independent experiments. Student’s t-test, p<0.001 (F) Wings were mounted, imaged, and measured after imaginal disc damage and regeneration in *osa*^*308*^/+ and wild-type (*w*^*1118*^) animals. n = 142 wings (*osa*^*308*^/+) and 284 wings (*w*^*1118*^) from three independent experiments. (G) Wild-type (*w*^*1118*^) adult wing after disc regeneration. Anterior is up. (H) *osa*^*308*^/+ adult wing after disc regeneration. Arrows show five anterior-specific markers in the posterior compartment: anterior crossveins (red), alula-like costa bristles (orange), margin vein (green), socketed bristles (blue), and change of wing shape with wider distal portion of the wing, similar to the anterior compartment (purple). (I) Quantification of the number of Posterior-to-Anterior transformation markers described in (H) in each wing after damage and regeneration of the disc, using wings that were 75% normal size or larger, comparing *osa*^*308*^/+ wings to wild-type (*w*^*1118*^) wings, n = 51 wings (*osa*^*308*^/+) and 45 wings (*w*^*1118*^), from 3 independent experiments. Chi-square test p < 0.001. Error bars are s.e.m. Scale bars are 500μm for all adult wings images. * p < 0.05, ** p < 0.01, ***p < 0.001 Student’s *t*-test.

Interestingly, the *osa*^*308*^/+ adult wings also showed severe patterning defects after damage and regeneration of the disc (Fig. 2G-I). Specifically, the posterior compartment of the *osa*^*308*^/+ wings had anterior features after wing pouch ablation, but had normal wings when no tissue damage was induced (Fig. S1B). To quantify the extent of the posterior-to-anterior (P-to-A) transformations, we quantified the number of anterior features in the posterior of each wing, including socketed bristles and ectopic veins on the posterior margin, an ectopic anterior crossvein (ACV), costal bristles on the alula, and an altered shape that has a narrower proximal and wider distal P compartment (Schuster and Smith-Bolton, 2015) (Fig. 2I). While *w*^*1118*^ adult wings that had regenerated as discs had a low level of P-to-A transformations, 75% of the *osa*^*308*^/+ wings had P-to-A transformations, and 83% of these transformed wings had 4 or 5 anterior markers in the posterior of the wing. Thus, Osa is required to preserve posterior cell fate during regeneration, suggesting that the BAP complex regulates cell fate after damage.

### Reducing the core SWI/SNF components to varying levels produces either the BAP or PBAP phenotype

Because mutants of the BAP or PBAP complex-specific components showed distinct phenotypes, we also screened mutants of the core components for regeneration phenotypes. Interestingly, mutants or RNAi lines that reduced levels of the core components were split between the two phenotypes. For example, *brm*^*2*^/+ discs and discs expressing a *Bap111* RNAi construct regenerated poorly, resulting in small wings (Fig. 3A,B), while *Bap55*^*LL05955*^/+ discs, *mor*^*1*^/+ discs, and discs expressing a *Bap60* RNAi construct regenerated to produce larger wings overall that showed P-to-A transformations (Table S1, Fig. 3C-G, Fig. S1A).

**Fig 3.**
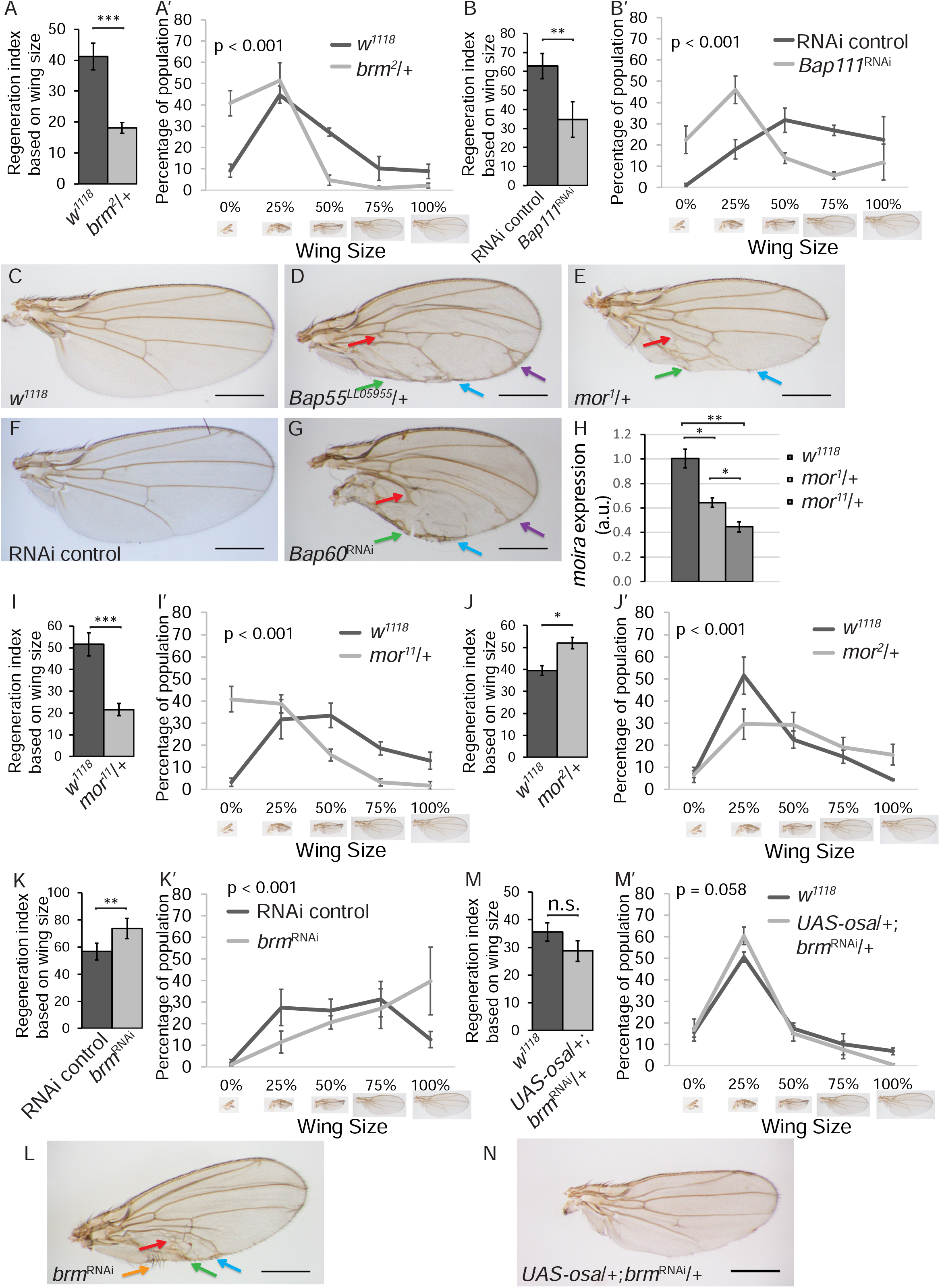
SWI/SNF core components are required for both growth and posterior fate during wing disc regeneration. (A) Comparison of the size of adult wings after imaginal disc damage and regeneration in *brm*^*2*^/+ and wild-type (*w*^*1118*^) animals. n = 142 wings (*brm*^*2*^/+) and 224 wings (*w*^*1118*^) from 3 independent experiments, student’s t-test p < 0.001. (A’) Chi-square test p < 0.001 across all wing sizes. (B) Comparison of the size of adult wings after imaginal disc damage and regeneration in animals expressing *Bap111* RNAi and control animals. n = 264 wings (*Bap111* RNAi) and 291 wings (control) from 3 independent experiments. The control for RNAi lines is VDRC 15293 in all experiments, student’s t-test p < 0.01. (B’) Chi-square test p < 0.001 across all wing sizes. (C-G) Adult wing after disc regeneration of wild-type (*w*^*1118*^) (C), *Bap55*^*LL05955*^/+ (D), *mor*^*1*^/+ (E), RNAi control (F) or *Bap60* RNAi (G). Anterior is up for all adult wing images. Arrows point to anterior features identified in the posterior compartment. Arrows show five anterior-specific markers in the posterior compartment: anterior crossveins (red), alula-like costa bristles (orange), margin vein (green), socketed bristles (blue), and change of wing shape with wider distal portion of the wing, similar to the anterior compartment (purple). (H) *moira* expression determined by qPCR of *mor*^*1*^/+, *mor*^*11*^/+ and wild-type (*w*^*1118*^) undamaged wing discs at R24. The graph shows fold change relative to wild-type (*w*^*1118*^) discs. (I) Comparison of the size of adult wings after imaginal disc damage and regeneration in *mor*^*11*^/+ and wild-type (*w*^*1118*^) animals. n = 114 wings (*mor*^*11*^/+) and 328 wings (*w*^*1118*^) from 3 independent experiments, student’s t-test p < 0.001. (I’) Chi-square test p < 0.001 across all wing sizes. (J) Comparison of the size of adult wings after imaginal disc damage and regeneration in *mor*^*2*^/+ and wild-type (*w*^*1118*^) animals. n = 134 wings (*mor*^*2*^/+) and 414 wings (*w*^*1118*^) from 3 independent experiments, student’s t-test p < 0.05. (J’) Chi-square test p < 0.001 across all wing sizes. (K) Comparison of the size of adult wings after imaginal disc damage and regeneration in animals expressing *brm* RNAi and control animals. n = 234 wings (*brm* RNAi) and 281 wings (control) from 3 independent experiments, student’s t-test p < 0.01. (K’) Chi-square test p < 0.001 across all wing sizes. (L) Adult wing after disc regeneration while expressing *brm* RNAi. (M) Comparison of the size of adult wings after imaginal disc damage and regeneration in *UAS-osa*/+*; brm* ^RNAi^/+ and wild-type (*w*^*1118*^) animals. n = 117 wings (*UAS-osa*/+*; brm* ^RNAi^/+) and 348 wings (*w*^*1118*^) from 3 independent experiments, student’s t-test not significant. (M’) Chi-square test across all wing sizes p = 0.058, not significant at α = 0.05 level. (N) Adult wing after imaginal disc regeneration in *UAS-osa*/+*; brm*^RNAi^/+ animal. Error bars are s.e.m. Scale bars are 500μm for all adult wing images. * p < 0.05, ** p < 0.01, ***p < 0.001 Student’s *t*-test.

Given that the SWI/SNF complexes require the function of the scaffold Mor and the ATPase Brm (Mashtalir et al., 2018; Moshkin et al., 2007), it was surprising that reduction of Mor showed the BAP phenotype while reduction of Brm showed the PBAP phenotype. However, it is likely that some of the mutants and RNAi lines caused stronger loss of function than others. A stronger reduction in function would result in malfunction of both BAP and PBAP, and show the reduced regeneration phenotype, masking any patterning defects. By contrast, a weaker reduction in function could mainly affect the BAP complex. For example, *Bap60* RNAi, which caused patterning defects after wing disc regeneration, only induced a moderate reduction in mRNA levels, suggesting that it causes a weak loss of function (Fig S1C). Although it is unclear why a weaker reduction of function would mainly affect the BAP complex, it is possible that the BAP complex is less abundant than the PBAP complex, such that a slight reduction in a core component would have a greater effect on the amount of BAP in the tissue. Therefore, we hypothesized that stronger or weaker loss of function of the same core complex component might show different phenotypes.

To test this hypothesis, we used a strong loss-of-function *mor* mutant, *mor*^*11*^ (gift from J. Kennison, Fig. S1D), and two hypomorphic *mor* mutants *mor*^*1*^ *and mor*^*2*^ (Kennison and Tamkun, 1988). Indeed, *mor*^*11*^*/+* undamaged wing discs had significantly less *mor* transcript than *mor*^*1*^*/+* or control undamaged wing discs (Fig. 3H). Interestingly, *mor*^*11*^*/+* animals showed the poor regeneration phenotype similar to the PBAP complex-specific *Bap170*^*Δ135*^/+ mutants (Fig. 3I), while *mor*^*1*^*/+* and *mor*^*2*^*/+* showed the enhanced regeneration phenotype and the P-to-A transformation phenotype similar to the BAP complex-specific *osa*^*308*^/+ mutants (Fig. 3E,J, S1Table). To confirm these findings we also used an amorphic allele of *brm* and an RNAi line that targets *brm* to reduce the levels of the core component *brm*. *brm*^*2*^ was generated through ethyl methanesulfonate mutagenesis and causes a loss of Brm protein (Elfring et al., 1998; Kennison and Tamkun, 1988). The *brm* RNAi causes a partial reduction in transcript, as *rn>brmRNAi* undamaged wing discs had less *brm* transcript than control undamaged wing discs (Fig. S1E). *brm*^*2*^/+ animals showed the small wing phenotype after disc damage, indicating poor regeneration (Fig. 3A). By contrast, knockdown of *brm* by expressing the *brm* RNAi construct during tissue ablation induced larger wings and P-to-A transformations (Fig. 3K,L). Thus, slight reduction of the core SWI/SNF components, through *mor*^*1*^, *brm* RNAi, or *Bap60* RNAi, produced the BAP phenotype, whereas stronger reduction of the core components, through *mor*^*11*^, produced the PBAP phenotype, suggesting that it is easier to compromise BAP function than to compromise PBAP function. If it is easier to compromise BAP function because there is less BAP complex in regenerating wing disc cells, overexpression of the BAP-specific component Osa would lead to an increase in the amount of BAP complex and rescue the *brm* RNAi phenotype. Indeed, overexpression of *osa* in regenerating tissue rescued the enhanced wing size and P-to-A transformations induced by *brm* RNAi (Fig. 3M,N).

### The PBAP complex is required for Myc upregulation and cell proliferation during regrowth

To identify when the defect in regrowth occurs in PBAP complex mutants, we measured the regenerating wing pouch using expression of the pouch marker *nubbin* in *w*^*1118*^ controls, *Bap170*^*Δ135*^/+ and *brm*^*2*^/+ mutants, as well as in the *osa*^*308*^/+ BAP mutant for comparison. The regenerating wing pouches of *Bap170*^*Δ135*^/+ mutant animals were not different in size compared to *w*^*1118*^ animals at 0, 12, or 24 hours after tissue damage (R0, R12 or R24). However, the *Bap170*^*Δ135*^/+ regenerating wing pouches were smaller than *w*^*1118*^ by 36 hours after tissue damage (R36), shortly before the *Bap170*^*Δ135*^/+ mutant animals pupariated and entered metamorphosis (Fig. 4A-C). *brm*^*2*^/+ mutant animals also had smaller regenerating wing pouches by R24 (Fig. S2A-C). By contrast, the regenerating *osa*^*308*^/+ wing pouches regrew at the same rate as controls (Fig. S2D-H). To determine whether the *Bap170*^*Δ135*^/+ mutant animals had a slower rate of pro-liferation during regeneration, we quantified the number of mitotic cells by immunostaining for phospho-histone H3 (PH3) in the regenerating wing pouch. A 35% decrease in the number of PH3-positive cells was observed in *Bap170*^*Δ135*^/+ mutants (Fig. 4D-F, Fig. S2I). While smaller adult wings could also be caused by increased cell death in the regenerating tissue, we did not find an increase in cell death in *Bap170*^*Δ135*^/+ regenerating wing discs as marked by immunostaining for cleaved caspase Dcp1 (Fig. S2J,K).

**Fig 4.**
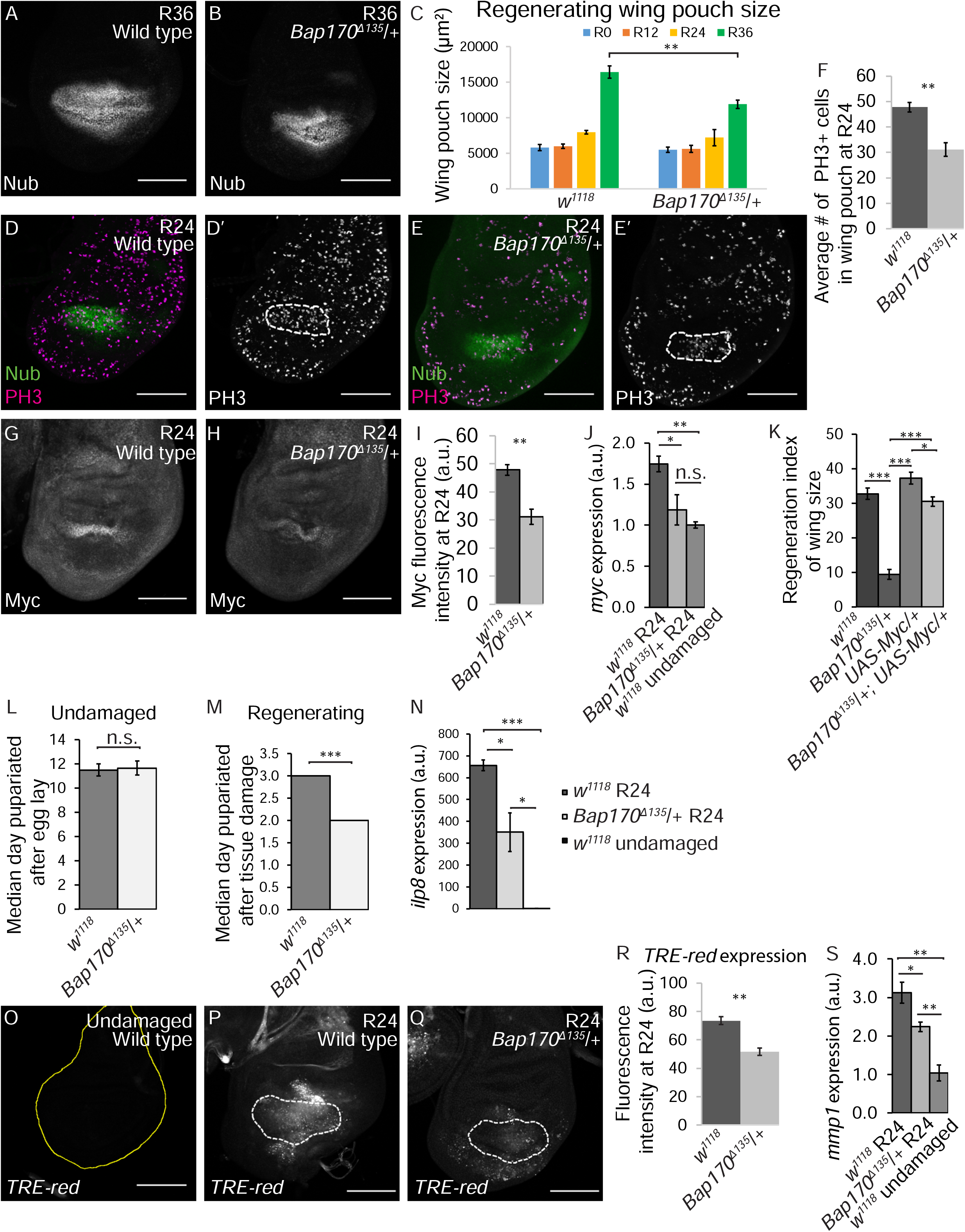
Decreased *Bap170* expression limits regenerative growth and pupariation delay. (A) Wild-type (*w*^*1118*^) regenerating wing disc at R36 with wing pouch marked by anti-Nubbin (green) immunostaining. (B) *Bap170*^*Δ135*^/+ regenerating wing disc at R36 with wing pouch marked by anti-Nubbin (green) immunostaining. (C) Comparison of regenerating wing pouch size at 0, 12, 24, and 36 hours after imaginal disc damage in *Bap170*^*Δ135*^/+ and wild-type (*w*^*1118*^) animals. (D-E) Regenerating wild-type (*w*^*1118*^) (D) and *Bap170*^*Δ135*^/+ (E) wing discs at R24 with Nubbin (green) and PH3 (magenta) immunostaining. Dashed white outline shows the regenerating wing primordium labeled with Nubbin. (F) Average number of mitotic cells (marked with PH3 immunostaining) in the wing primordium (marked by anti-Nubbin) at R24 in *Bap170*^*Δ135*^/+ and wild-type (*w*^*1118*^) animals. n = 8 wing discs (*Bap170*^*Δ135*^/+) and 10 wing discs (*w*^*1118*^). (G-H) Wild-type (*w*^*1118*^) (G) and *Bap170*^*Δ135*^/+ (H) regenerating wing discs at R24 with Myc immunostaining. (I) Quantification of anti-Myc immunostaining fluorescence intensity in the wing pouch in *Bap170*^*Δ135*^/+ and wild-type (*w*^*1118*^) regenerating wing discs at R24. n = 9 wing discs (*Bap170*^*Δ135*^/+) and 9 wing discs (*w*^*1118*^). (L) Median time to pupariation for animals during normal development at 18°C. n = 103 pupae (*Bap170*^*Δ135*^/+) and 227 pupae (*w*^*1118*^) from 3 independent experiments. Student’s *t*-test not significant. (M) Median time to pupariation for animals after tissue damage (30°C) and regeneration (18°C). n = 117 pupae (*Bap170*^*Δ135*^/+) and 231 pupae (*w*^*1118*^) from 3 independent experiments. Because the temperature shift to 30°C in the ablation protocol increases the developmental rate, the pupariation timing of regenerating animals (M) cannot be compared to the undamaged control animals (L). student’s t-test p<0.001. (N) *ilp8* expression examined by qPCR of *Bap170*^*Δ135*^/+ and wild-type (*w*^*1118*^) regenerating wing discs at R24. The graph shows fold change relative to wild-type (*w*^*1118*^) undamaged discs. (O-Q) Expression of *TRE-Red*, a JNK signaling reporter, in wild-type (*w*^*1118*^) undamaged (O), as well as wild-type (*w*^*1118*^) (P) and *Bap170*^*Δ135*^/+ (Q) regenerating wing discs at R24. Yellow outline shows the wing disc in (O). White dashed lines show the wing pouch in (P) and (Q) as marked by anti-Nub. (R) Quantification of *TRE-Red* fluorescence intensity in *Bap170*^*Δ135*^/+ and wild-type (*w*^*1118*^) regenerating wing pouches at R24. n = 12 wing discs (*Bap170*^*Δ135*^/+) and 14 wing discs (*w*^*1118*^). (S) *mmp1* expression examined by qPCR of wild-type (*w*^*1118*^) and *Bap170*^*Δ135*^/+ regenerating wing discs at R24, and wild-type (*w*^*1118*^) undamaged discs. The graph shows fold change relative to wild-type (*w*^*1118*^) regenerating discs at R24. Scale bars are 100μm for all wing discs images. * p < 0.05, ** p < 0.01, *** p < 0.001, Student’s *t*-test.

To identify why proliferation was reduced in *Bap170*^*Δ135*^/+ mutants, we examined levels of Myc, an important growth regulator that is upregulated during *Drosophila* wing disc regeneration (Smith-Bolton et al., 2009). In mammals, *c-myc* is a direct target of the SWI/SNF BAF complex, which is similar to *Drosophila* BAP (Nagl et al., 2006), but a role for the PBAP complex in regulating the *Drosophila Myc* gene has not been established. Myc protein levels were significantly reduced in *Bap170*^*Δ135*^/+ and *brm*^*2*^/+ regenerating wing pouches compared to wild-type regenerating wing pouches (Fig. 4G-I and Fig. S3A-D). Myc transcriptional levels were also significantly lower in *Bap170*^*Δ135*^/+ regenerating wing discs compared to wild-type regenerating discs (Fig. 4J). By contrast, there was no change in Myc levels in *osa*^*308*^/+ mutants (Fig. S3E-G), indicating that PBAP, but not BAP, is required for upregulation of Myc after tissue damage. To determine the extent to which reduction of Myc expression was responsible for the poor regeneration phenotype in BAP complex mutants, we overexpressed Myc in the *Bap170*^*Δ135*^/+ background during regeneration. Indeed, the *Bap170*^*Δ135*^/+, *UAS-Myc*/+ animals regenerated similar to the *w*^*1118*^ controls and significantly better than *Bap170*^*Δ135*^/+ animals, demonstrating partial rescue of the poor regeneration phenotype (Fig. 4K and Fig. S3H).

### The PBAP complex is required for the delay in pupariation induced by tissue damage

Damaged imaginal discs delay pupariation by expressing the peptide ILP8, which delays the production of ecdysone and onset of metamorphosis, providing more time for damaged tissue to regenerate (Colombani et al., 2012; Garelli et al., 2012). To determine whether the SWI/SNF complexes regulate the timing of metamorphosis, we quantified the pupariation rate in *w*^*1118*^ and *Bap170*^*Δ135*^/+ regenerating animals, and identified the day on which 50% of the larvae had pupariated. Without tissue damage, *Bap170*^*Δ135*^/+ mutants pupariated slightly later than *w*^*1118*^ animals (Fig. 4L and Fig. S4A), but the difference is not significant. However, after wing disc damage, more than half of the *Bap170*^*Δ135*^/+ mutant animals had pupariated by 2 days after damage, whereas more than half of the *w*^*1118*^ animals had not pupariated until 3 days after damage, giving the mutants 1/3 less time to regenerate (Fig. 4M and Fig. S4B). To uncover why *Bap170*^*Δ135*^/+ animals had less regeneration time, we quantified *ilp8* transcript levels. Indeed, *Bap170*^*Δ135*^/+ animals had about 50% less *ilp8* mRNA (Fig. 4N), suggesting that the PBAP complex is required for *ilp8* expression.

### The PBAP complex regulates expression of JNK signaling targets

SWI/SNF complexes can be recruited by transcription factors to act as co-activators of gene expression (Becker and Workman, 2013). Regenerative growth and the pupariation delay are regulated by JNK signaling (Bergantinos et al., 2010; Bosch et al., 2008; Colombani et al., 2012; Garelli et al., 2012; Skinner et al., 2015). Thus, it is possible that PBAP is recruited to JNK signaling targets like *ilp8* by the AP-1 transcription factor, which acts downstream of JNK (Perkins et al., 1988), and that PBAP is required for full activation of these targets. To determine whether *Bap170* is required for JNK-dependent transcription, we examined the activity of the TRE-Red reporter, which is comprised of four AP-1 binding sites (TREs) driving the expression of a DsRed.T4 reporter gene (Chatterjee and Bohmann, 2012) in *w*^*1118*^ and *Bap170*^*Δ135*^/+ regenerating wing discs. The TRE-Red intensity was significantly decreased in the *Bap170*^*Δ135*^/+ regenerating tissue compared to the *w*^*1118*^ regenerating tissue (Fig. 4O-R), indicating that PBAP is required for full activation of this AP-1 transcriptional activity reporter, similar to its requirement for expression of *ilp8*. Furthermore, expression of the JNK signaling target *mmp1* was significantly reduced in *Bap170*^*Δ135*^/+ regenerating wing discs at both the mRNA and protein levels (Fig. 4S and Fig. S4C-E). Thus, the PBAP complex plays a crucial role in activation of JNK signaling targets.

### The BAP complex maintains posterior cell fate during regeneration

After damage and regeneration of the disc, adult wings of *osa*^*308*^/+, *Bap55*^*LL05955*^/+, *mor*^*1*^/+, and *mor*^*2*^/+ discs, as well as discs expressing a *brm* RNAi construct or a *Bap60* RNAi construct, had anterior bristles and veins in the posterior compartment (Fig. 3C-G,K), but not after normal development (Fig. S1A, S1F-H). To identify when the P-to-A transformations occurred, we examined the expression of anterior- and posterior-specific genes during tissue regeneration. *engrailed* (*en*) is essential for posterior cell fate both in development and regeneration (Kornberg et al., 1985; Schuster and Smith-Bolton, 2015). To assess ability to maintain posterior cell fate, regenerating wing discs were dissected at different times during recovery (R) and immunostained for the posterior selector gene *en*. At 72 hours after damage (R72), in *osa*^*308*^/+ regenerating discs, *en* was expressed in some of the posterior compartment, but lost in patches (Fig. 5A-C). In addition, the proneural protein Acheate (Ac), which is expressed in sensory organ precursors in the anterior of wing discs (Skeath and Carroll, 1991), was ectopically expressed in the posterior (Fig. 5D-F) marking precursors to the ectopic socketed bristles found in the posterior of the adult wings. The anterior genes *cubitus interruptus* (*ci*) (Eaton and Kornberg, 1990) and *patched* (*ptc*) (Phillips et al., 1990) were also ectopically expressed in the posterior of the *osa*^*308*^/+ R72 regenerating wing discs (Fig. 6A-C). The ectopic expression of these anterior genes was not observed at R48, suggesting that the P-to-A fate transformations happened late during regeneration (Fig. S4F,G). Similarly, at R72, 80% of the *brm* RNAi wing discs had ectopic expression of the anterior genes *ptc* and *ci* in the posterior of the discs, while no expression of *ptc* or *ci* was observed in the posterior of control R72 discs (Fig. 6D,E).

**Fig 5.**
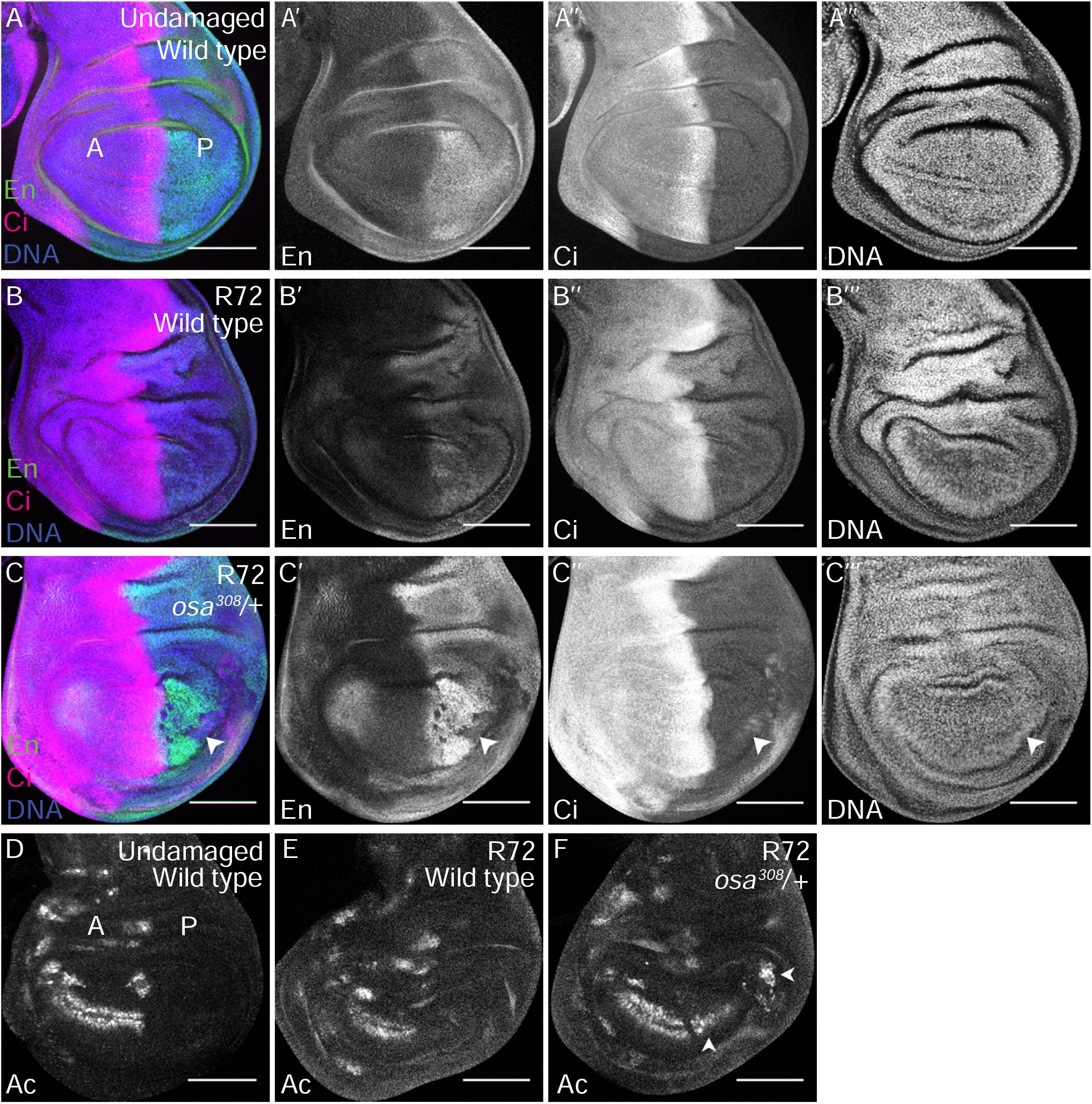
Reduction of Osa causes Posterior-to-Anterior transformations during wing disc regeneration. (A) Wild-type (*w*^*1118*^) undamaged wing disc with En (green) (A’) and Ci (magenta) (A’’) immunostaining. DNA (blue) (A’’’) was detected with Topro3 here and in subsequent panels. Anterior is left for all wing disc images. (B) Wild-type (*w*^*1118*^) regenerating wing disc at R72 with En (green) (B’) and Ci (magenta) (B’’) immunostaining and DNA (blue) (B’’’). (C) *osa*^*308*^/+ regenerating wing disc at R72 with En (green) (C’) and Ci (magenta) (C’’) immunostaining, and DNA (blue) (C’’’). Arrowhead points to the low En expression region in which Ci is expressed in the posterior compartment. (D) Wild-type (*w*^*1118*^) undamaged wing disc with Ac immunostaining. (E) Wild-type (*w*^*1118*^) regenerating wing disc at R72 with Ac immunostaining. (F) *osa*^*308*^/+ regenerating wing disc at R72 with Ac immunostaining. Arrowheads show Ac expression in the posterior compartment. Scale bars are 100μm for all wing discs images.

**Fig 6.**
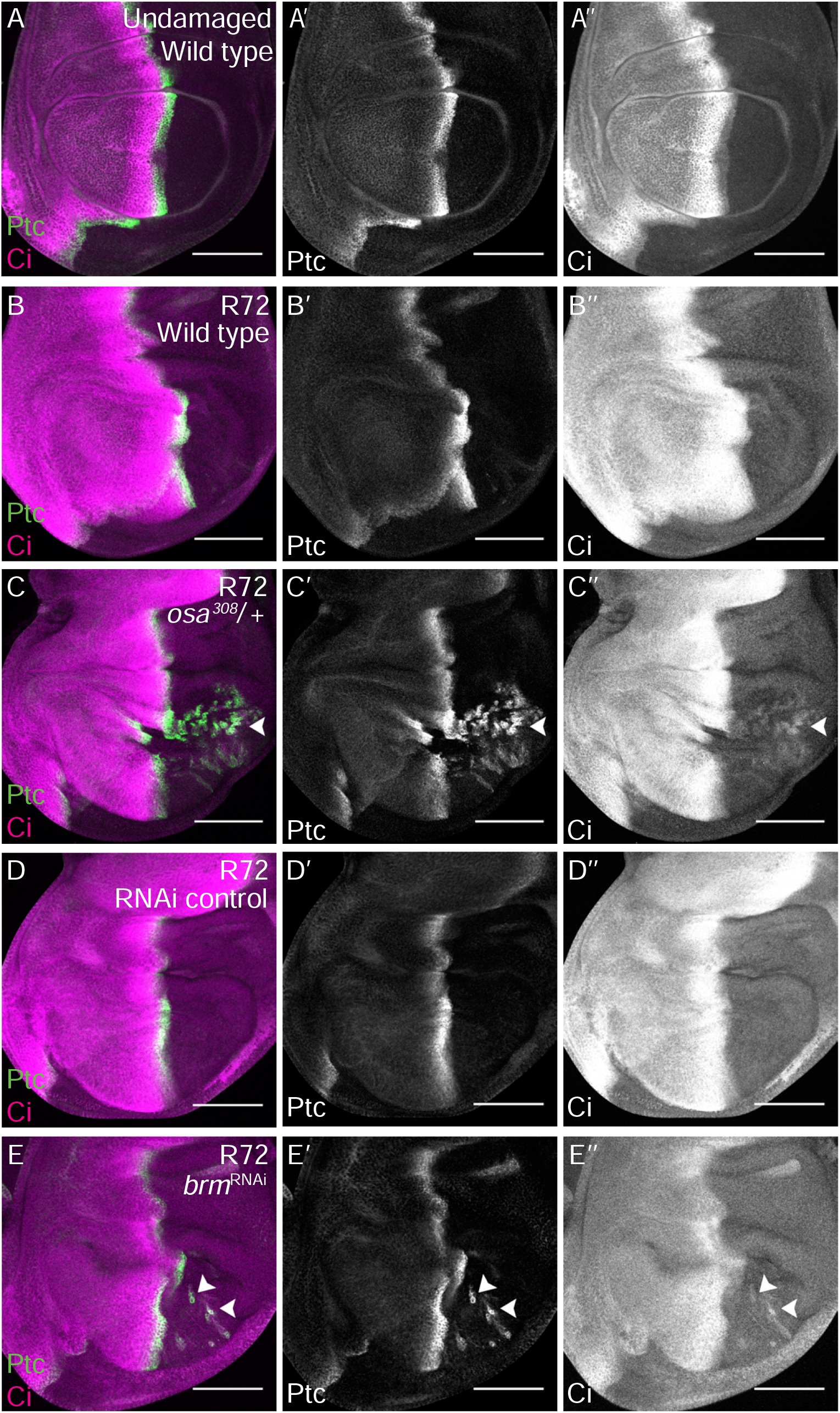
The BAP complex is required to maintain posterior cell fate during wing disc regeneration. (A) Wild-type (*w*^*1118*^) undamaged wing disc with Ptc (green) (A’) and Ci (magenta) (A’’) immunostaining. (B) Wild-type (*w*^*1118*^) regenerating wing disc at R72 with Ptc (green) (B’) and Ci (magenta) (B’’) immunostaining. (C) *osa*^*308*^/+ regenerating wing disc at R72 with Ptc (green) (C’) and Ci (magenta) (C’’) immunostaining. Arrowhead shows Ptc and Ci co-expression in the posterior compartment. (D) RNAi control regenerating wing disc at R72 with Ptc (green) (D’) and Ci (magenta) (D’’) immunostaining. (E) Regenerating wing disc of animals expressing *brm* RNAi at R72 with Ptc (green) (E’) and Ci (magenta) (E’’) immunostaining. Arrowheads show Ptc and Ci co-expression in the posterior compartment. Scale bars are 100μm for all wing disc images.

We previously showed that in *Drosophila* wing disc regeneration, elevated JNK increases expression of *en*, leading to PRC2-mediated silencing of the *en* locus in patches, and transformation of the *en*-silenced cells to anterior fate, and that Taranis prevents this misregulation of *en* and resulting P-to-A cell fate transformations (Schuster and Smith-Bolton, 2015). Thus, we wondered whether the BAP complex preserved *en* expression and posterior fate by reducing JNK signaling, or regulating *tara* expression, or working in parallel to Tara during the later stages of regeneration.

### The BAP complex does not regulate JNK signaling

To determine whether the BAP complex regulates JNK signaling, we examined the JNK reporter TRE-Red in *osa*^*308*^/+ and *w*^*1118*^ regenerating wing discs. In contrast to *Bap170*^*Δ135*^/+ mutants (Fig. 4O-R), TRE-Red intensity was not different between *osa*^*308*^/+ and *w*^*1118*^ regenerating tissue (Fig. 7A-C). Thus, the BAP complex acts to protect posterior cell fate downstream of or in parallel to JNK signaling.

**Fig 7.**
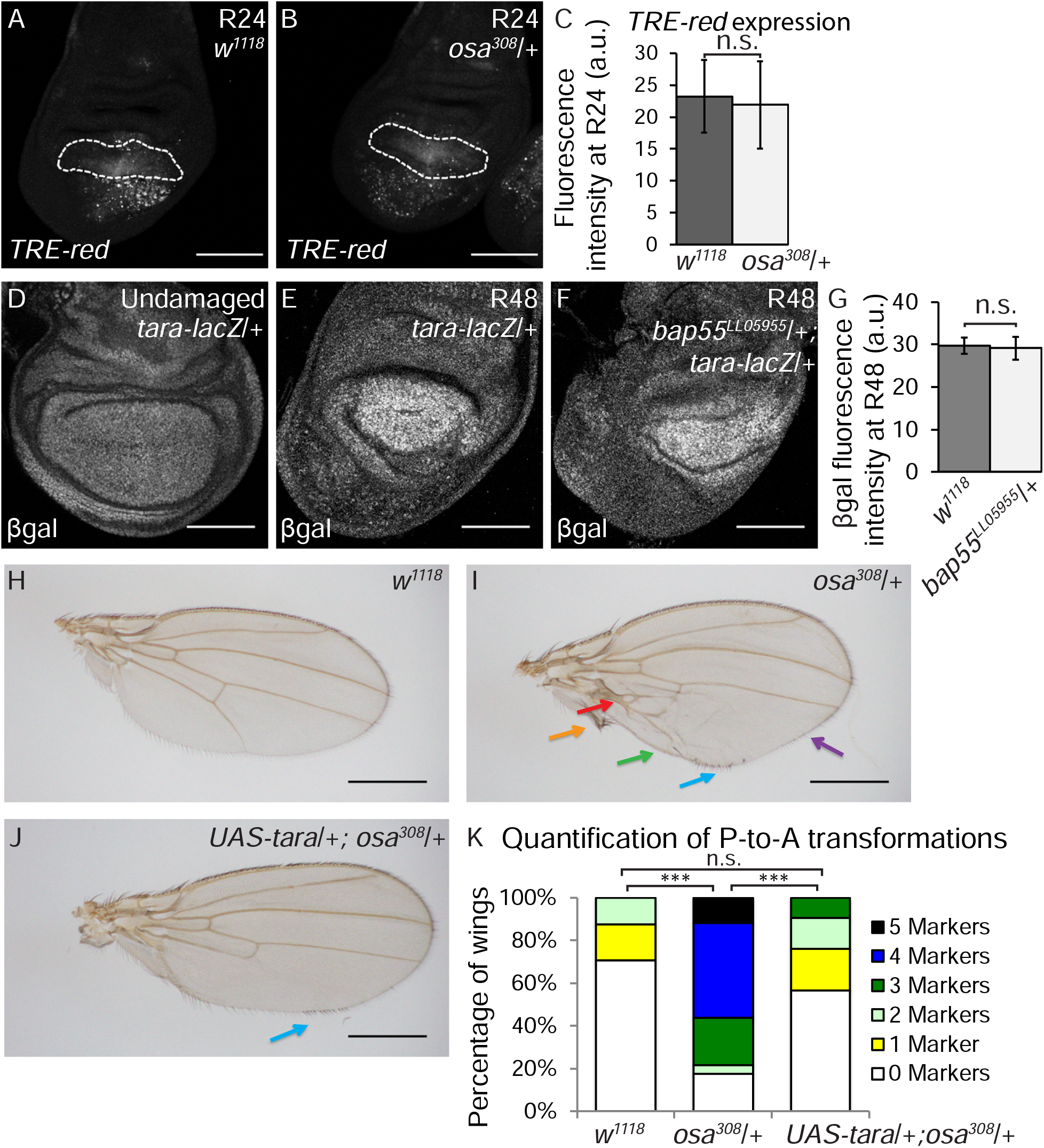
The BAP complex functions in parallel to Tara to prevent P-to-A transformations. (A-B) Expression of *TRE-Red*, a JNK signaling reporter, in wild-type (*w*^*1118*^) (A) and *osa*^*308*^/+ (B) regenerating wing discs at R24. Dashed white outline shows the regenerating wing primordium as marked by anti-Nub and excluding the debris field. (C) Quantification of *TRE-Red* expression fluorescence intensity in *osa*^*308*^/+ and wild-type (*w*^*1118*^) regenerating wing pouches at R24. n = 26 wing discs (*osa*^*308*^/+) and 31 wing discs (*w*^*1118*^). Error bars are s.e.m. (D-F) Tara expression detected with anti-β-gal immunostaining in *tara-lacZ*/+ undamaged (D), *tara-lacZ*/+ R48 (E) and *Bap55*^*LL05955*^/+; *tara-lacZ*/+ R48 (F) regenerating wing discs. (G) Quantification of β-gal expression via fluorescence intensity to determine levels of *tara-lacZ* expression in *Bap55*^*LL05955*^/+ and wild-type (*w*^*1118*^) regenerating wing pouches at R48. n = 8 wing discs (*Bap55*^*LL05955*^/+) and 9 wing discs (*w*^*1118*^). Error bars are s.e.m. (H-J) Adult wings after disc regeneration in wild-type (*w*^*1118*^) (H), *osa*^*308*^/+ (I) and *UAS-tara*/+; *osa*^*308*^/+ (J) animals. Arrows show five anterior-specific markers in the posterior compartment: anterior crossveins (red), alula-like costa bristles (orange), margin vein (green), socketed bristles (blue), and change of wing shape with wider distal portion of the wing, similar to the anterior compartment (purple). Anterior is up for all adult wing images. (K) Quantification of the number of Posterior-to-Anterior transformation markers described above in each wing after damage and regeneration of the disc, comparing *UAS-tara*/+; *osa*^*308*^/+ wings to *osa*^*308*^/+ and wild-type (*w*^*1118*^) wings, n = 21 wings (*UAS-tara*/+; *osa*^*308*^/+), n = 16 wings (*osa*^*308*^/+) and n = 34 wings (*w*^*1118*^), from 3 independent experiments. *** p < 0.001, Chi-square test. Chi-square test measuring *UAS-tara*/+; *osa*^*308*^/+ against *w*^*1118*^, p = 0.86, is not significant. Scale bars are 100μm for all wing discs images. Scale bars are 500μm for all adult wings images. * p < 0.05, ** p < 0.01, Student’s *t*-test for (C) and (G).

### The BAP complex functions in parallel to Taranis to preserve cell fate

Because *tara* is regulated transcriptionally after tissue damage (Schuster and Smith-Bolton, 2015), we examined whether the BAP complex is required for *tara* upregulation in the regenerating tissue. Using a *tara-lacZ* enhancer trap, we assessed expression in *Bap55*^*LL05955*^/+ regenerating wing discs, which had the same P-to-A transformations as the *osa*^*308*^/+ regenerating discs. No change in *tara-lacZ* expression was identified in the regenerating wing pouches, (Fig. 7D-G), indicating that the damage-dependent *tara* expression was not downstream of BAP activity.

To determine whether Tara can suppress the P-to-A transformations induced by the reduction of BAP, we overexpressed Tara using *UAS-tara* under control of *rn-Gal4* in the *osa*^*308*^/+ mutant animals, generating elevated Tara levels in the *rn*-expressing cells that survived the tissue ablation. Indeed, the P-to-A transformation phenotype in *osa*^*308*^/+ mutant animals was rescued by Tara overexpression (Fig. 7H-K). To rule out the possibility that Tara regulates *osa* expression, we quantified Osa immunostaining in *tara/+* mutant regenerating tissue. Osa protein levels did not change during regeneration, and were unchanged in *tara*^*1*^/+ mutant regenerating discs (Fig. S4H-M). Taken together, these data indicate that the BAP complex likely functions in parallel to Tara to constrain *en* expression, preventing auto-regulation and silencing of *en*, thereby protecting cell fate from changes induced by JNK signaling during regeneration.

### The enhanced growth in BAP mutants is caused by ectopic AP boundaries

The increased wing size after disc regeneration in *tara/+* animals was due to loss of *en* in patches of cells, which generated aberrant juxtaposition of anterior and posterior tissue within the posterior compartment. These ectopic AP boundaries established ectopic Dpp morphogen gradients (Schuster and Smith-Bolton, 2015), which can stimulate extra growth in the posterior compartment (Tanimoto et al., 2000). To determine whether the *osa*/+ regenerating discs also had ectopic AP boundaries and ectopic morphogen gradients, we immunostained for Ptc to mark AP boundaries and phospho-Smad to visualize gradients of Dpp signaling. Indeed, Ectopic regions of Ptc expression were surrounded by ectopic pSmad gradients in osa^308^/+ regenerating discs (Fig. 8A-C). Thus, the enhanced regeneration in *osa*^*308*^/+ and other SWI/SNF mutant animals was likely a secondary result of the patterning defect. Furthermore, pupariation occurred later in *osa*^*308*^*/+* regenerating animals compared to *w*^*1118*^ regenerating animals (Fig. S4N,O), which provided more time for regeneration in the mutants. Such a delay in pupariation can by caused by aberrant proliferation (Colombani et al., 2012; Garelli et al., 2012) in addition to tissue damage, and the combination of the two likely led to the increase in delay in metamorphosis seen specifically in mutants with P-to-A transformations.

**Fig 8.**
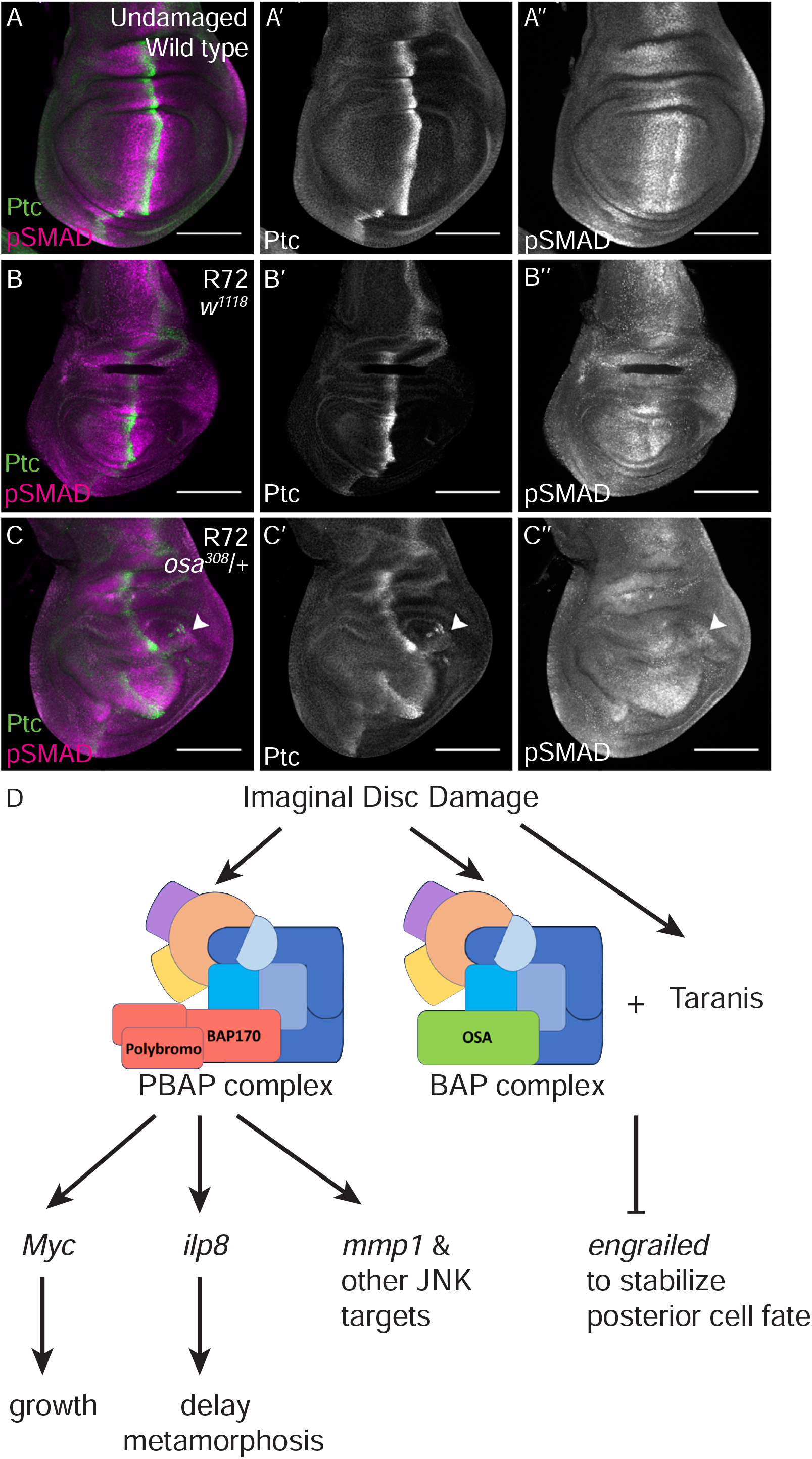
Cell fate changes induce ectopic AP boundaries in the posterior compartment during wing disc regeneration. (A) Wild-type (*w*^*1118*^) undamaged wing disc with Ptc (green) (A’) and pSMAD (magenta) (A’’) immunostaining. (B) Wild-type (*w*^*1118*^) regenerating wing disc at R48 with Ptc (green) (B’) and pSMAD (magenta) (B’’) immunostaining. (C) *osa*^*308*^/+ regenerating wing disc at R48 with Ptc (green) (C’) and Ci (magenta) (C’’) immunostaining. (D) Proposed working model for the functions of the PBAP and BAP complexes in regeneration.

## Discussion

To address the question of how regeneration genes are regulated in response to tissue damage, we screened a collection of mutants and RNAi lines that affect a significant number of the chromatin regulators in *Drosophila*. Most of these mutants had regeneration phenotypes, confirming that these genes are important for both promoting and constraining regeneration and likely facilitate the shift from the normal developmental program to the regeneration program, and back again. The variation in regeneration phenotypes among different chromatin regulators and among components of the same multi-unit complexes supports our previous finding that damage activates expression of genes that both promote and constrain regeneration (Khan et al., 2017). Such regulators of regeneration may be differentially affected by distinct mutations that affect the same chromatin-modifying complexes, resulting in different phenotypes.

We have demonstrated that both *Drosophila* SWI/SNF complexes play essential but distinct roles during epithelial regeneration, controlling multiple aspects of the process, including growth, developmental timing, and cell fate (Fig. 8D). Furthermore, our work has identified multiple likely targets, including *mmp1, myc*, *ilp8*, and *en*. Indeed, analysis of data from a recent study that identified regions of the genome that transition to open chromatin after imaginal disc damage showed such damage-responsive regions near *Myc*, *mmp1*, and *ilp8* (Vizcaya-Molina et al., 2018). While previous work has suggested that chromatin modifiers can regulate regeneration (Blanco et al., 2010; Fukuda et al., 2012; Jin et al., 2013; Jin et al., 2015; Pfefferli et al., 2014; Scimone et al., 2010; Skinner et al., 2015; Stewart et al., 2009; Sun et al., 2016; Tseng et al., 2011; Wang et al., 2008; Xiong et al., 2013), and that the chromatin near *Drosophila* regeneration genes is modified after damage (Harris et al., 2016; Vizcaya-Molina et al., 2018), our results suggest that these damage-responsive loci are not all coordinately regulated in the same manner. The SWI/SNF complexes target different subsets of genes, and it will not be surprising if different cofactors or transcription factors recruit different complexes to other subsets of regeneration genes.

Is the requirement for the SWI/SNF complexes for growth and conservation of cell fate in the wing disc specific to regeneration? In contrast to *tara,* which is required for posterior wing fate only after damage and regeneration (Schuster and Smith-Bolton, 2015), loss of *mor* in homozygous clones during wing disc development caused loss of *en* expression in the posterior compartment (Brizuela and Kennison, 1997), although this result was interpreted to mean that *mor* promotes rather than constrains *en* expression, which is the opposite of our observations. Importantly, undamaged *mor* heterozygous mutant animals did not show patterning defects (Fig. S1G,H), while damaged heterozygous mutant animals did (Fig. 3E), indicating that regenerating tissue is more sensitive to reductions in SWI/SNF levels than normally developing tissue. Furthermore, *osa* is required for normal wing growth (Terriente-Félix and de Celis, 2009), but reduction of *osa* levels did not compromise growth during regeneration (Fig. 2D). Thus, while some functions of SWI/SNF during regeneration may be the same as during development, other functions of SWI/SNF are unique to regeneration.

SWI/SNF complexes help organisms respond rapidly to stressful conditions or changes in the environment. For example, SWI/SNF is recruited by the transcription factor DAF-16/FOXO to promote stress resistance in *Caenorhabditis elegans* (Riedel et al., 2013), and the *Drosophila* BAP complex is required for the activation of target genes of the NF-κB signaling transcription factor Relish in immune responses (Bonnay et al., 2014). Here we show that the *Drosophila* PBAP complex is similarly required after tissue damage for activation of target genes of the JNK signaling transcription factor AP-1 after tissue damage. Interestingly, the BAF60a subunit, a mammalian homolog of *Drosophila* BAP60, directly binds the AP-1 transcription factor and stimulates the DNA-binding activity of AP-1 (Ito et al., 2001), suggesting that this role may be conserved.

In summary, we have demonstrated that the two SWI/SNF complexes regulate different aspects of wing imaginal disc regeneration, implying that activation of the regeneration program is controlled by changes in chromatin, but that the mechanism of regulation is likely different for subsets of regeneration genes. Future identification of all genes targeted by BAP and PBAP after tissue damage, the factors that recruit these chromatin-remodeling complexes, and the changes they induce at these loci will deepen our understanding of how unexpected or stressful conditions lead to rapid activation of the appropriate genes.

## Acknowledgements

The authors would like to thank A. Brock and K. Schuster for critical reading of the manuscript and helpful discussions; A. Dingwall, D. Bohmann, J. Kennison, J. Treisman, M Cleary, S. Cohen, the Bloomington Drosophila Stock Center (NIH P40OD018537), the Vienna *Drosophila* Resource Center, and the Developmental Studies Hybridoma Bank for fly stocks and reagents.

