## Supplemental materials for "The *Drosophila* SWI/SNF chromatin-remodeling complexes BAP and PBAP play separate roles in regulating growth and cell fate during regeneration"

**Table S1. Screen of Chromatin Regulators.** For each line, the regeneration index was calculated by summing the product of approximate wing size (0%, 25%, 50%, 75% and 100%) and the corresponding percentage of wings for each wing size. The $\Delta$ Index, which is the difference between the regeneration indices of the line being tested and the control tested simultaneously, was calculated by subtracting the regeneration index of the control from the regeneration index of the mutant or RNAi line. A cutoff $\Delta$ index of 10% was set, over which we considered the regenerative capacity to be affected. Green indicates lines that had a higher regeneration index compared to the control, purple indicates lines that had a lower regeneration index compared to the control. * = $\Delta$ Index ≥ 10 or ≤ -10, ** = $\Delta$ Index ≥ 20 or ≤ -20, *** = $\Delta$ Index ≥ 30 or ≤ -30.

**Fig S1. Roles of the SWI/SNF components in regeneration and development**

(A) Average Δ regeneration index of chromatin regulator mutants and RNAi lines screen. The Δ regeneration Index is the difference between the regeneration indices of the line being tested and the control tested simultaneously. Δ regeneration index was calculated as described in materials and methods.

(B) Undamaged adult wings of $osa^{308}$/+ animals.

(C) *Bap60* expression examined by qPCR of $Bap60^{RNAi}$ and control undamaged wing discs. The graph shows fold change relative to control discs. RNAi lines were crossed to $w^{1118}$; +; *rn-GAL4, tubGAL80*$^{ts}$*/TM6B* and kept at 18°C. Temperature shift to 30°C at day 7 for 24 hours then back to 18°C. Wing discs of non-Tubby larvae were dissected at 24 hours after shifting back to 18°C.

(D) Complementation test for $mor^{1}$ and $mor^{11}$ mutants.

(E) *brm* expression examined by qPCR of $brm^{RNAi}$ and control undamaged wing discs. The graph shows fold change relative to control discs. RNAi lines were crossed to $w^{1118}$; +; *rn-GAL4, tubGAL80*$^{ts}$*/TM6B* and kept at 18°C. Temperature shift to 30°C at day 7 for 24 hours then back to 18°C. Wing discs of non-Tubby larvae were dissected at 24 hours after shifting back to 18°C.

(F-H) Undamaged adult wings of $Bap55^{LL05955}$/+ (E), $mor^{1}$/+ (F) and $mor^{2}$/+ (G) animals.

Error bars are S.E.M.. Scale bars are 500μm for all adult wings images. * $p < 0.05$, ** $p < 0.01$, *** $p < 0.001$ Student's *t*-test for (C) and (E).

**Fig S2. The PBAP complex is required for regenerative growth whereas the BAP complex is not.**

(A) Wild-type (*w*$^{1118}$) regenerating wing disc at R24 with wing pouch marked by anti-Nubbin immunostaining.

(B) *brm*$^{2}$/+ regenerating wing disc at R24 with wing pouch marked by anti-Nubbin immunostaining.

(C) Comparison of regenerating wing pouch size at 24 hours after imaginal disc damage in *brm*$^{2}$/+ and wild-type (*w*$^{1118}$) animals. n = 11 wing discs (*brm*$^{2}$/+) and 10 wing discs (*w*$^{1118}$).

(D) Wild-type (*w*$^{1118}$) regenerating wing disc at R24 with wing pouch marked by anti-Nubbinimmunostaining.

(E) *osa*$^{308}$/+ regenerating wing disc at R24 with wing pouch marked by anti-Nubbin immunostaining.

(F) Wild-type (*w*$^{1118}$) regenerating wing disc at R48 with wing pouch marked by anti-Nubbin immunostaining.

(G) *osa*$^{308}$/+ regenerating wing disc at R48 with wing pouch marked by anti-Nubbin immunostaining.

(H) Comparison of regenerating wing pouch size at 24 and 48 hours after imaginal disc damage and regeneration in *osa*$^{308}$/+ and wild-type (*w*$^{1118}$) animals. At R24, n = 26 wing discs (*osa*$^{308}$/+) and 27 wing discs (*w*$^{1118}$). At R48, n = 6 wing discs (*osa*$^{308}$/+) and 21 wing discs (*w*$^{1118}$).

(I) Average number of mitotic cells (marked with PH3 immunostaining) per µm$^{2}$ in the regenerating wing primordium at R24 in *Bap170*$^{\Delta 135}$/+ and wild-type (*w*$^{1118}$) animals. n = 8 wing discs (*Bap170*$^{\Delta 135}$/+) and 10 wing discs (*w*$^{1118}$).

(J) Wild-type (*$w^{1118}$*) regenerating wing disc at R24 with Nubbin (green) (J') and cleaved caspase Dcp1 (magenta)(J'') immunostaining marking the debris field, and DNA (blue) was detected with Topro3 here in subsequent panel. (J''').

(K) *$bap170^{\Delta 135}$*/+ regenerating wing disc at R24 with Nubbin (green)(K') and cleaved caspase Dcp1 (magenta)(K'') immunostaining, and DNA (blue)(K''').

Error bars are S.E.M.. Scale bars are 100μm for all wing discs images. * $p < 0.05$, ** $p < 0.01$, Student's *t*-test.

**Fig S3. The PBAP complex regulates Myc in regeneration whereas the BAP complex does not.**

(A) Wild-type (*w*$^{1118}$) undamaged wing disc with Myc (magenta) (A') and Nubbin (green) (A'') immunostaining. DNA (blue) (A''') was detected with Topro3.

(B-C) Wild-type (*w*$^{1118}$) (B) and *brm*$^{2}$/+ (C) regenerating wing discs at R24 with Myc immunostaining.

(D) Quantification of anti-Myc immunostaining fluorescence intensity in the wing pouch in *brm*$^{2}$/+ and wild-type (*w*$^{1118}$) regenerating wing discs at R24. n = 11 wing discs (*brm*$^{2}$/+) and 12 wing discs (*w*$^{1118}$).

(E-F) Wild-type (*w*$^{1118}$) (E) and *osa*$^{308}$/+ (F) regenerating wing discs at R24 with Myc immunostaining.

(G) Quantification of anti-Myc immunostaining fluorescence intensity in the wing pouch in *osa*$^{308}$/+ and wild-type (*w*$^{1118}$) regenerating wing discs at R24. n = 28 wing discs (*osa*$^{308}$/+) and 27 wing discs (*w*$^{1118}$).

(H) Comparison of the size of adult wings after imaginal disc damage and regeneration in wild-type (*w*$^{1118}$), *Bap170*$^{\Delta 135}$/+, *Bap170*$^{\Delta 135}$/+*; UAS-Myc*/+, and *UAS-Myc*/+ animals. n = 364 wings (*w*$^{1118}$), 194 wings (*Bap170*$^{\Delta 135}$/+), 194 wings (*Bap170*$^{\Delta 135}$/+*; UAS-Myc*/+) and 294 wings (*UAS-Myc*/+) from 3 independent experiments. Chi-square test for *Bap170*$^{\Delta 135}$/+ and *Bap170*$^{\Delta 135}$/+*; UAS-Myc*/+ has $p < 0.001$

Error bars are S.E.M.. Scale bars are 100µm for all wing discs images. * $p < 0.05$, ** $p < 0.01$, Student's *t*-test for (D) and (G).

**Fig S4. The function of BAP and PBAP in regeneration and development**

(A) Pupariation rates of animals during normal development at 18°C. n = 103 pupae (*Bap170*$^{\Delta 135}$/+) and 227 pupae (*w*$^{1118}$) from 3 independent experiments. Student's *t*-test not significant.

(B) Pupariation rates of animals after tissue damage (30°C) and regeneration (18°C). n = 117 pupae (*Bap170*$^{\Delta 135}$/+) and 231 pupae (*w*$^{1118}$) from 3 independent experiments. Because the temperature shift to 30°C in the ablation protocol increases the developmental rate, the pupariation timing of regenerating animals (B) cannot be compared to the undamaged control animals (A). Chi-square test $p < 0.001$.

(C-E) *mmp1* expression examined by immunofluorescence in wild-type (*w*$^{1118}$) (C) and *Bap170*$^{\Delta 135}$/+ (D) regenerating wing discs at R24. Quantification in (E). n= 19 (*w*$^{1118}$) and 17 (*Bap170*$^{\Delta 135}$/+), error bars are S.E.M., p=0.00041.

(F) Wild-type (*w*$^{1118}$) regenerating wing disc at R48 with Ptc (green)(F') and Ci (magenta)(F'') immunostaining.

(G) *osa*$^{308}$/+ regenerating wing disc at R48 with Ptc (green)(G') and Ci (magenta)(G'') immunostaining.

(H-K) Wild-type (*w*$^{1118}$) regenerating wing discs at 0, 24, 48, and 72 hours after imaginal disc damage with Osa immunostaining.

(L-M) Wild-type (*w*$^{1118}$) (L) and *tara*$^{1}$/+ (M) regenerating wing discs at R48 with Osa immunostaining.

(N) Pupariation rates of animals during normal development at 18°C. n = 79 pupae (*osa*$^{308}$/+) and 173 pupae (*w*$^{1118}$) from 3 independent experiments. Chi-square test $p < 0.001$, student's t-test at day 11 $p<0.01$.

(O) Pupariation rates of animals after tissue damage (30°C) and regeneration (18°C). n = 101 pupae (*osa*$^{308}$/+) and 155 pupae (*w*$^{1118}$) from 3 independent experiments. Chi-square test $p < 0.001$, student's t-test at day 3 $p<0.01$. Because the temperature shift to 30°C in the ablation protocol increases the developmental rate, the pupariation timing of regenerating animals (O) cannot be compared to the undamaged control animals (N).

Scale bars are 100μm for all wing discs images. Scale bars are 100μm for all wing discs images. Error bars are S.E.M. except where noted. * $p < 0.05$, ** $p < 0.01$, ***<0.001 for Student's *t*-test.

S1 Table. Screen of chromatin regulators

| Allele or RNAi | Complex | Regeneration Phenotype | ∆ Index | Phenotype Strength |
|---|---|---|---|---|
| *Pc*$^{3}$ | PRC1 | | 7% | |
| *Psc*$^{1}$ | PRC1 | | 9% | |
| *Psc*$^{e24}$ | | **Reduced** | **-20%** | ** |
| *Sce*$^{1}$ | PRC1 | **Enhanced** | **18%** | * |
| *Scm*$^{D1}$ | PRC1 | **Enhanced** | **46%** | *** |
| *E(z)*$^{731}$ | PRC2 | **Reduced** | **-14%** | * |
| *Su(z)12*$^{2}$ | PRC2 | | -9% | |
| *esc*$^{21}$ | PRC2 | **Enhanced** | **26%** | ** |
| *Caf1-55*$^{DG25308}$ | PRC2, NURF | **Reduced** | **-19%** | * |
| *escl*$^{d01514}$ | PRC2 | **Reduced** | **-20%** | ** |
| *phol*$^{81A}$ | PhoRC | **Enhanced** | **41%** | *** |
| *ash2*$^{1}$ | COMPASS, COMPASS-like | **Enhanced** | **16%** | * |
| *Utx*$^{f01321}$ | COMPASS-like | **Enhanced** | **16%** | * |
| *ash1*$^{22}$ | ASH1 | | 7% | |
| *E(bx)*$^{nurf301\text{-}3}$ | NURF | **Reduced** | **-17%** | * |
| *Nurf-38*$^{k16102}$ | NURF | | -1% | |
| *Mi-2*$^{4}$ | NuRD | | 5% | |
| *brm*$^{2}$ | SWI/SNF (BAP & PBAP) | **Reduced** | **-23%** | ** |
| *brm*$^{RNAi\ VDRC37721}$ | | **Enhanced** | **18%** | * |
| *osa*$^{308}$ | SWI/SNF(BAP) | **Enhanced** | **28%** | ** |
| *Bap170*$^{\Delta 135}$ | SWI/SNF(PBAP) | **Reduced** | **-19%** | * |
| *polybromo*$^{\Delta 86}$ | SWI/SNF(PBAP) | **Reduced** | **-20%** | ** |
| *Snr1*$^{E2}$ | SWI/SNF (BAP & PBAP) | | 5% | |
| *Snr1*$^{SR21}$ | | **Enhanced** | **17%** | * |
| *mor*$^{1}$ | SWI/SNF (BAP & PBAP) | **Enhanced** | **11%** | * |
| *mor*$^{2}$ | | **Enhanced** | **12%** | * |
| *mor*$^{11}$ | | **Reduced** | **-30%** | *** |
| *mor*$^{12}$ | | **Enhanced** | **13%** | * |
| *mor*$^{RNAi\ VDRC6969}$ | | **Enhanced** | **42%** | *** |
| *Bap55*$^{LL05955}$ | SWI/SNF (BAP & PBAP) | **Enhanced** | **23%** | ** |
| *Bap60*$^{RNAi\ VDRC12673}$ | SWI/SNF (BAP & PBAP) | **Enhanced** | **12%** | * |
| *Bap111*$^{RNAi\ VDRC104361}$ | SWI/SNF (BAP & PBAP) | **Reduced** | **-28%** | ** |
| *psq*$^{E39}$ | GBP | **Enhanced** | **15%** | * |

| | | | | |
|---|---|---|---|---|
| *Rbf*$^{14}$ | dREAM | **Enhanced** | **22%** | ** |
| *Dsp1*$^{EP355}$ | | **Enhanced** | **25%** | ** |
| *grh*$^{IM}$ | | | 6% | |
| *lolal*$^{K02512}$ | | | 1% | |
| *Pcl*$^{5}$ | | **Enhanced** | **16%** | * |
| *HDAC1*$^{def24}$ | Sin3A/HDAC complex | **Enhanced** | **20%** | ** |
| *Sirt1*$^{2A\text{-}7\text{-}11}$ | SIRT1-LSD1 complex | **Enhanced** | **23%** | ** |
| *vtd*$^{4}$ | cohesin | **Enhanced** | **47%** | *** |
| *Su(z)2*$^{1.b7}$ | | **Reduced** | **-14%** | * |
| *gpp*$^{03342}$ | | **Reduced** | **-14%** | * |
| *mod(mdg4)*$^{L3101}$ | | **Enhanced** | **19%** | * |
| *su(Hw)*$^{e04061}$ | gypsy chromatin insulator complex | **Enhanced** | **25%** | ** |
| *lid*$^{10424}$ | | **Enhanced** | **23%** | ** |
| *Asx*$^{XF23}$ | | **Enhanced** | **11%** | * |
| *dom*$^{LL05537}$ | TIP60 complex | | -3% | |
| *E(Pc)*$^{1}$ | TIP60 complex | **Enhanced** | **41%** | *** |
| *kis*$^{1}$ | | | 0% | |
| *kto*$^{1}$ | mediator | **Reduced** | **-11%** | * |
| *skd*$^{2}$ | mediator | **Enhanced** | **21%** | ** |

Fig S1

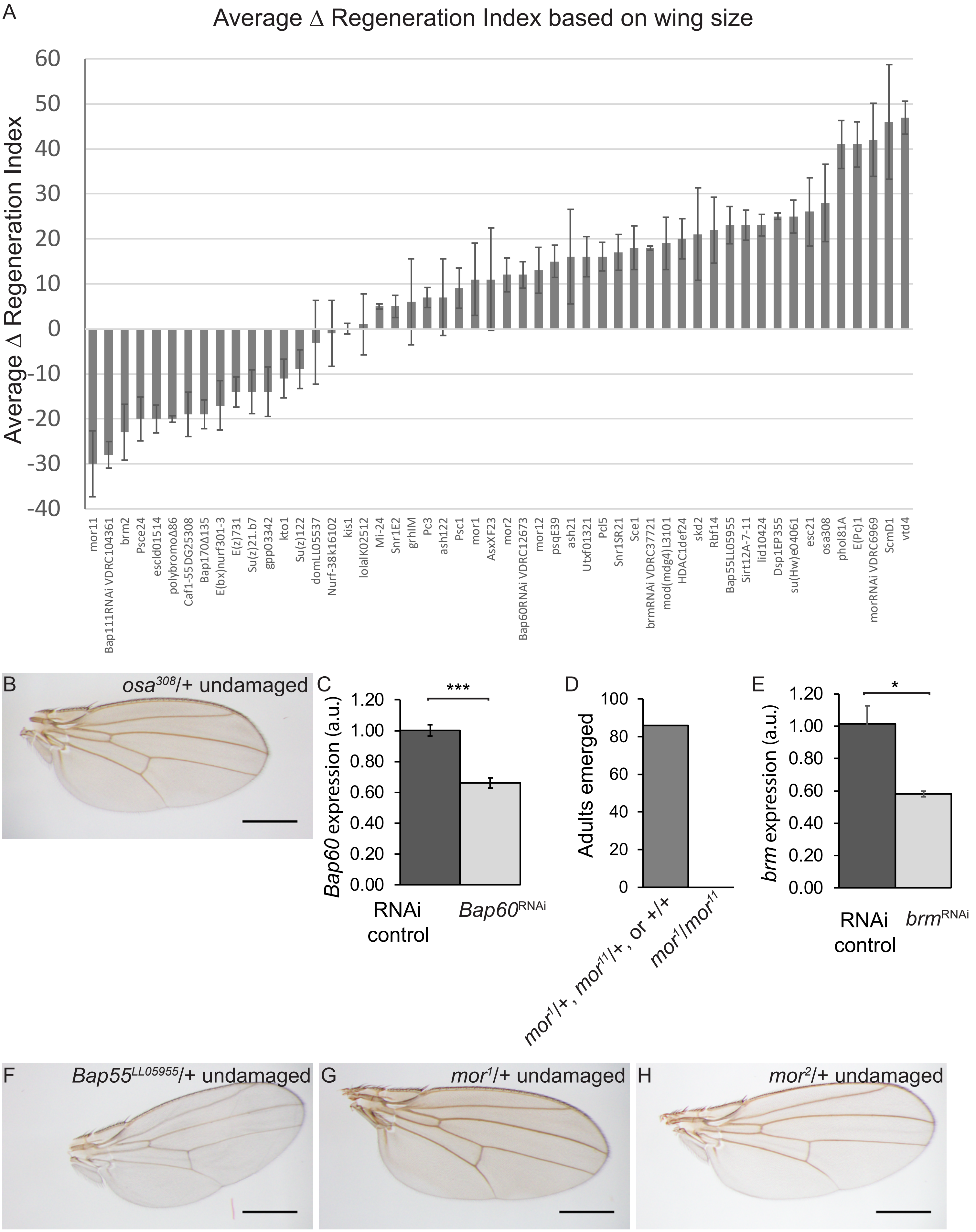

A
Average ∆ Regeneration Index based on wing size
Average ∆ Regeneration Index
60
50
40
30
20
10
0
-10
-20
-30
-40
mor11
Bap111RNAi VDRC104361
brm2
Psce24
escld01514
polybromoΔ86
Caf1-55DG25308
Bap170Δ135
E(bx)nurf301-3
E(z)731
Su(z)21.b7
gpp03342
kto1
Su(z)122
domLL05537
Nurf-38k16102
kis1
lolalK02512
Mi-24
Snr1E2
grhIM
Pc3
ash122
Psc1
mor1
AsxXF23
mor2
Bap60RNAi VDRC12673
mor12
psqE39
ash21
Utxf01321
Pcl5
Snr1SR21
Sce1
brmRNAi VDRC37721
mod(mdg4)L3101
HDAC1def24
skd2
Rbf14
Bap55LL05955
Sirt12A-7-11
lid10424
Dsp1EP355
su(Hw)e04061
esc21
osa308
phol81A
E(Pc)1
morRNAi VDRC6969
ScmD1
vtd4
B
osa308/+ undamaged
C
***
Bap60 expression (a.u.)
1.20
1.00
0.80
0.60
0.40
0.20
0.00
RNAi control
Bap60RNAi
D
Adults emerged
100
80
60
40
20
0
mor1/+, mor11/+, or +/+
mor1/mor11
E
*
brm expression (a.u.)
1.20
1.00
0.80
0.60
0.40
0.20
0.00
RNAi control
brmRNAi
F
Bap55LL05955/+ undamaged
G
mor1/+ undamaged
H
mor2/+ undamaged

Fig S2

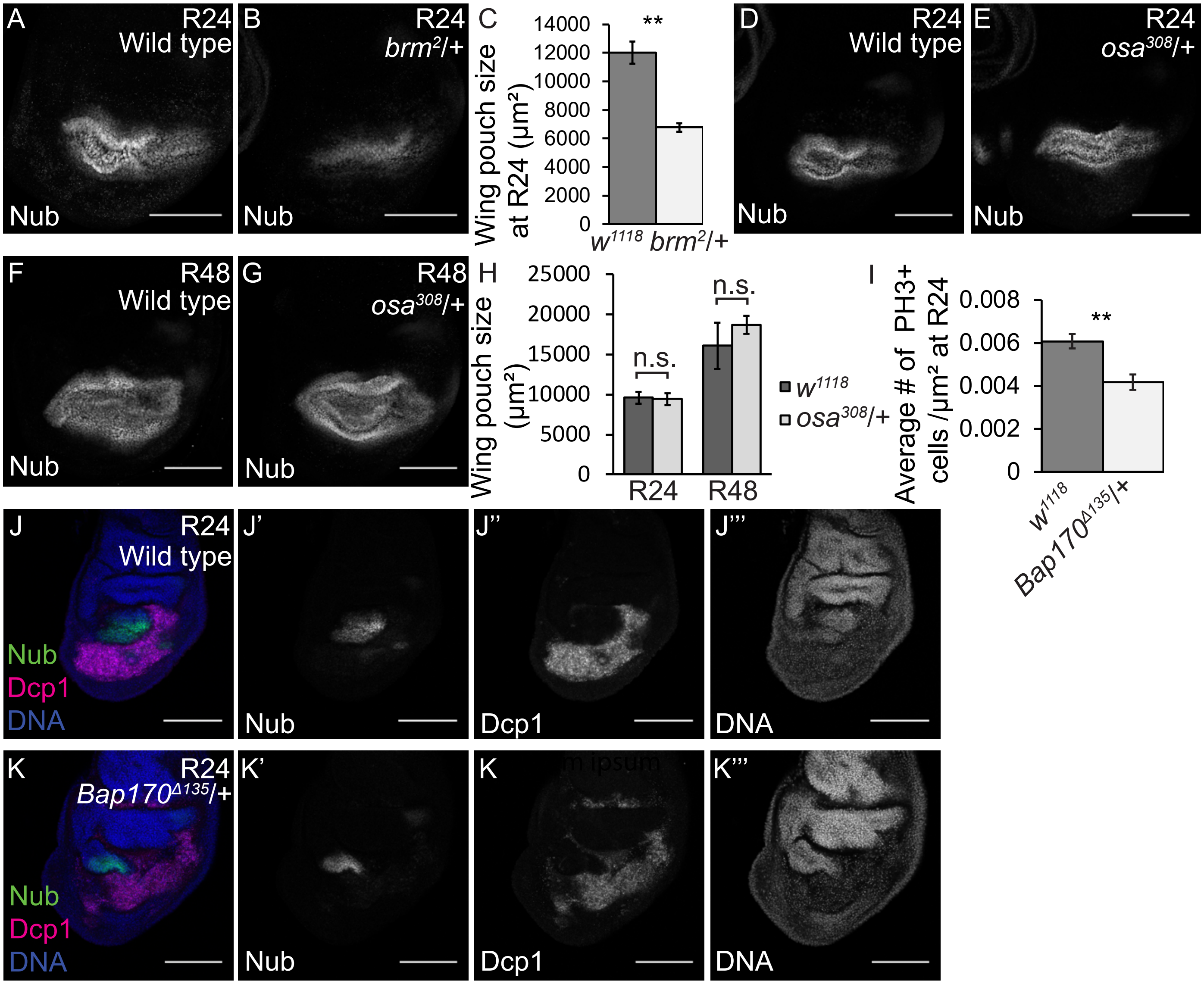
A
R24
Wild type
Nub
B
R24
brm2/+
Nub
C
Wing pouch size at R24 (μm²)
14000
12000
10000
8000
6000
4000
2000
0
**
w1118
brm2/+
D
R24
Wild type
Nub
E
R24
osa308/+
Nub
F
R48
Wild type
Nub
G
R48
osa308/+
Nub
H
Wing pouch size (μm²)
25000
20000
15000
10000
5000
0
n.s.
n.s.
R24
R48
w1118
osa308/+
I
Average # of PH3+ cells /μm² at R24
0.008
0.006
0.004
0.002
0
**
w1118
Bap170Δ135/+
J
R24
Wild type
Nub
Dcp1
DNA
J'
Nub
J''
Dcp1
J'''
DNA
K
R24
Bap170Δ135/+
Nub
Dcp1
DNA
K'
Nub
K
Dcp1
K'''
DNA

Fig S3

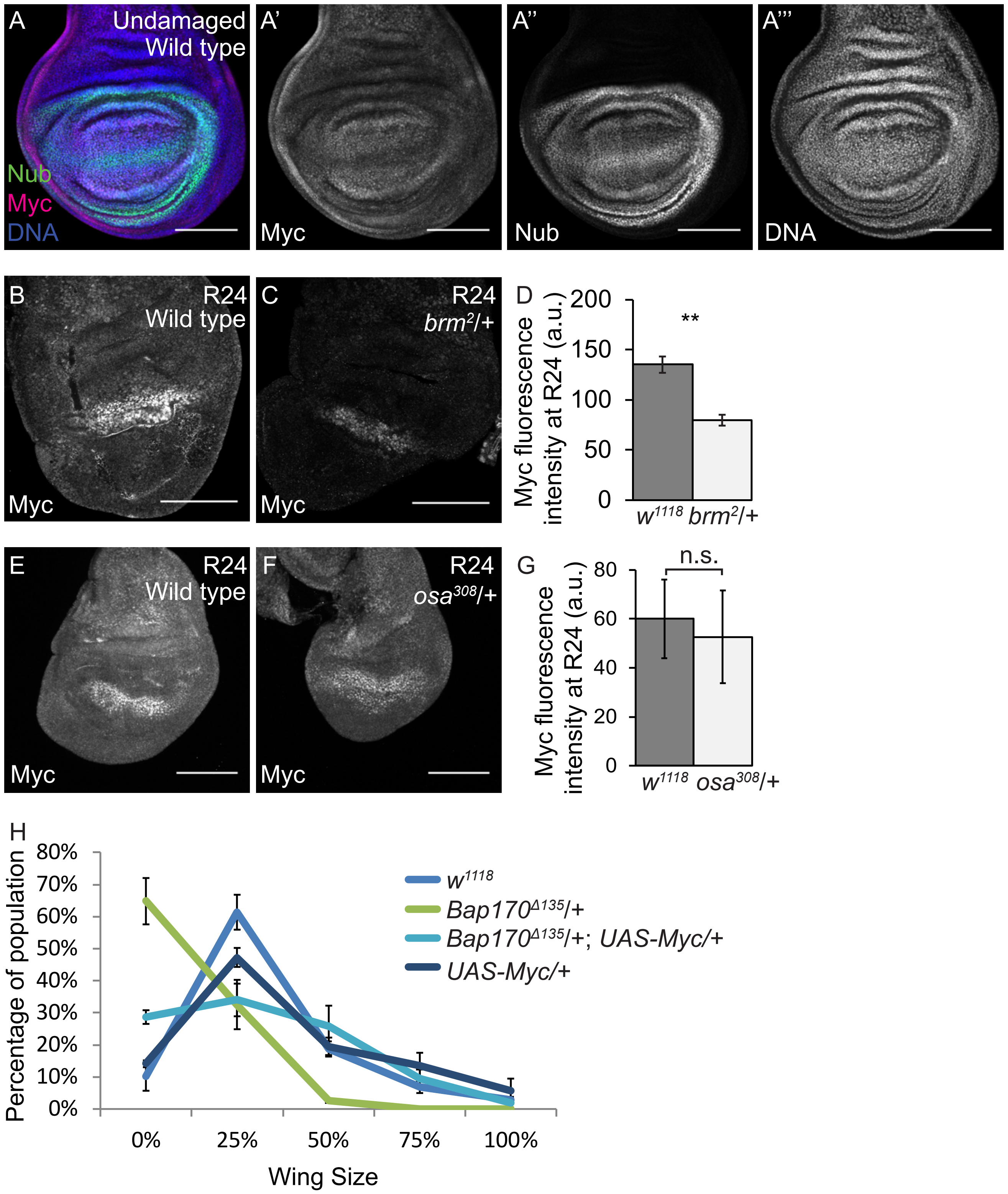
A
Undamaged
Wild type
Nub
Myc
DNA
A'
Myc
A''
Nub
A'''
DNA
B
R24
Wild type
Myc
C
R24
brm2/+
Myc
D
Myc fluorescence intensity at R24 (a.u.)
200
150
100
50
0
**
w1118
brm2/+
E
R24
Wild type
Myc
F
R24
osa308/+
Myc
G
Myc fluorescence intensity at R24 (a.u.)
80
60
40
20
0
n.s.
w1118
osa308/+
H
Percentage of population
80%
70%
60%
50%
40%
30%
20%
10%
0%
0%
25%
50%
75%
100%
Wing Size
w1118
Bap170Δ135/+
Bap170Δ135/+; UAS-Myc/+
UAS-Myc/+

Fig S4

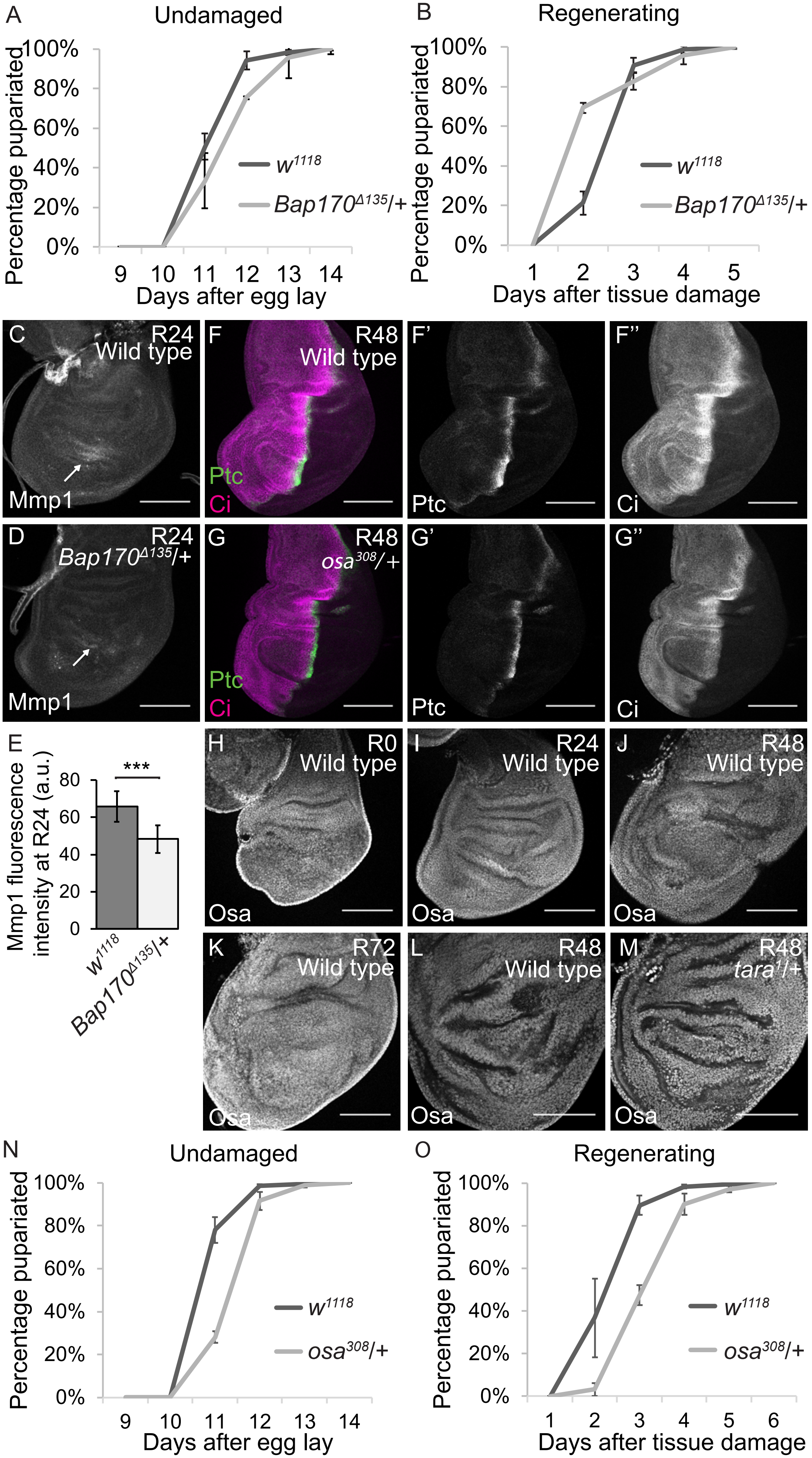

A
Undamaged
Percentage pupariated
100%
80%
60%
40%
20%
0%
9 10 11 12 13 14
Days after egg lay
w1118
Bap170Δ135/+
B
Regenerating
Percentage pupariated
1 2 3 4 5
Days after tissue damage
w1118
Bap170Δ135/+
C
R24
Wild type
Mmp1
D
R24
Bap170Δ135/+
Mmp1
E
Mmp1 fluorescence intensity at R24 (a.u.)
***
80
60
40
20
0
w1118
Bap170Δ135/+
F
R48
Wild type
Ptc
Ci
F'
Ptc
F''
Ci
G
R48
osa308/+
Ptc
Ci
G'
Ptc
G''
Ci
H
R0
Wild type
Osa
I
R24
Wild type
Osa
J
R48
Wild type
Osa
K
R72
Wild type
Osa
L
R48
Wild type
Osa
M
R48
tara1/+
Osa
N
Undamaged
Percentage pupariated
9 10 11 12 13 14
Days after egg lay
w1118
osa308/+
O
Regenerating
Percentage pupariated
1 2 3 4 5 6
Days after tissue damage
w1118
osa308/+
